# Native Mass Spectrometry Analysis of Cullin RING Ubiquitin E3 Ligase Complexes in the Context of Targeted Protein Degradation

**DOI:** 10.64898/2025.12.04.692288

**Authors:** Louise M Sternicki, Charlotte Crowe, Lianne H. E. Wieske, Alessio Ciulli, Sally-Ann Poulsen

## Abstract

The binding of PROTACs to their partner ubiquitin E3 ligase (E3) and a protein of interest (POI) is critical for PROTAC development and validation. Characterisation of PROTAC complexes by cryo-electron microscopy and X-ray crystallography is not always feasible, especially where species may be transient and protein structures may not resolve due to flexible domains or intrinsically disordered regions. More routine biophysical methods with broader applicability to varied samples is desirable to support the rapidly expanding targeted protein degradation field. The majority of PROTACs in development and in the clinic act through a Cullin RING E3 Ligase (CRL) of which the pentameric von Hippel-Lindau (VHL) Cullin 2 RING E3 complex (CRL2^VHL^) is the foundational example. Native mass spectrometry (nMS) can be used to characterise protein complexes but has not previously been used to characterise a full E3 or any E3-E2 interactions. Here, we show that CRL2^VHL^ is amenable to characterisation by nMS and its interactions with the other protein components integral to the targeted protein degradation mechanism can be observed. Specifically, we characterise binary, ternary and higher order complexes that comprise CRL2^VHL^, including the multiprotein systems of POI-PROTAC-CRL2^VHL^, CRL2^VHL^-E2-Ub, and POI-PROTAC-CRL2^VHL^-E2-Ub, all of which are essential in facilitating productive POI ubiquitination and degradation. We benchmarked the nMS with two POI examples (BRD4^BD2^ and KRAS) with their respective PROTACs (MZ1 and ACBI3) and were able to observe all relevant complexes across both systems. We anticipate that our findings will open avenues for nMS to integrate as an alternative experimental method enabling scalable characterisation of the intricate high mass multiprotein interactions central to the PROTAC mechanism of action.

## Introduction

The ubiquitin–proteasome system (UPS) is responsible for cellular protein degradation. It acts by covalently attaching ubiquitin (Ub), a 76 amino acid protein, to a lysine residue of the substrate protein via a three-enzyme cascade with a ubiquitin-activating enzyme (E1), a ubiquitin-conjugating enzyme (E2) and a ubiquitin ligase (E3). The E3 catalyses the transfer of Ub from the E2 to the target substrate, which then directs the substrate to the proteasome for degradation, **Figure 1**.^1^ It is possible to manipulate the UPS with small molecules called degraders that are designed to simultaneously bind an E3 and a protein of interest (POI), **Figure 1**.^2–4^ When applied for biological or therapeutic interventions this approach, known as targeted protein degradation, provides an almost unlimited scope to degrade any cellular protein, and degrader development is a rapidly expanding field.^2–5^ There are two well established categories of degrader molecules - PROTACs (PROteolysis TArgeting Chimeras) and molecular glues. They are distinctly different in architecture as PROTACs comprise two linked parts; the first part is a pharmacophore that binds the POI and the second part is a pharmacophore that binds to an E3, while molecular glues are made up from a single moiety that recognises both the POI and E3. Formation of a POI-degrader-E3 ternary complex followed by coordination to an E2 loaded with Ub (E2-Ub) is essential for ubiquitination.^6^ Despite the importance of the full POI-degrader-E3-E2-Ub complex for the proximity and orientation of the POI relative to the donor Ub, there are few structural studies that have captured this species. Furthermore, workflows to capture the multiprotein species of ubiquitination with native proteins at scale are lacking. This limits routine direct assessment of structural and mechanistic features of degrader-mediated protein complexes relevant to a degrader’s pharmacological activity.^7^

**Figure 1.**
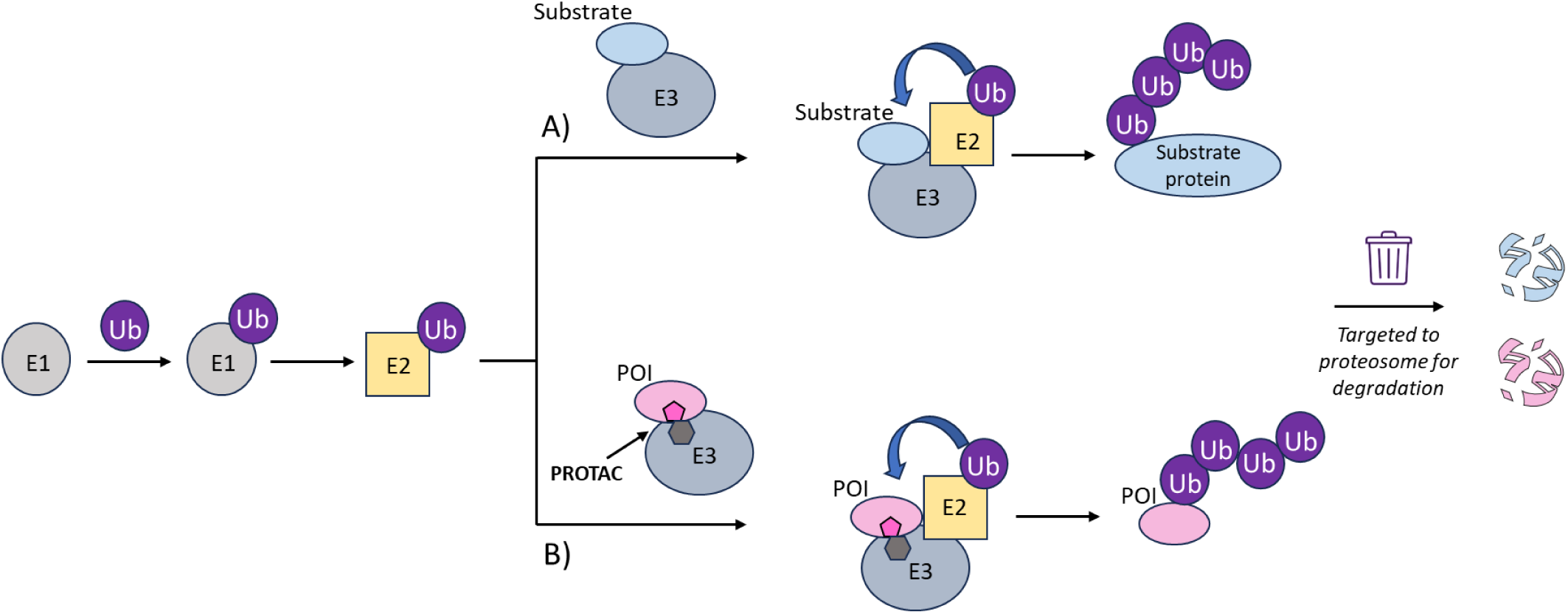
Comparison of native and PROTAC-hijacked protein degradation. Schematic overview of A) the ubiquitin–proteasome system (UPS) three-enzyme cascade with a ubiquitin-activating enzyme (E1), a ubiquitin-conjugating enzyme (E2) and a ubiquitin ligase (E3), and B) the same UPS pathways as hijacked by PROTAC molecules to induce ubiquitination and subsequent degradation of proteins of interest (POI) as non-native substrates.

There are >600 human E3 ligases with the Cullin RING ligase (CRL) E3 subfamily the most abundant.^8, 9^ CRLs are modular multiprotein complexes, comprising a Cullin scaffold protein (Cul1 – Cul5, Cul7 or Cul9), adaptor proteins (e.g. SKP1, EloB, EloC and/or DDB1), a RING-box protein (Rbx1 or Rbx2) that recruits the E2 enzyme loaded with Ub and a substrate-receptor protein that recruits the substrate protein for eventual degradation.^1, 8, 9^ Most reported structural studies of POI-degrader-E3 ternary complexes utilise the minimal E3 ligase VCB, this is a trimeric species comprising VHL (Von Hippel-Lindau protein) as the substrate-receptor protein with EloB (elongin B) and EloC (elongin C) as adapter proteins. Most notably VCB is without a Cullin scaffold protein or RING-box protein so cannot interact with E2-Ub.^10^ Although robust (VCB is readily expressed and has excellent stability) there are drawbacks to this minimal approach as the measured ternary complex binding is relatively far removed from the downstream biological activity sought in degrader development. The focus of this manuscript is native mass spectrometry (nMS) analysis of CRL2^VHL^, the foundational full E3 ligase that has been central in development of functional degraders. CRL2^VHL^ is a pentameric protein complex comprising VCB (a VHL-EloB-EloC trimer), Cul2 and Rbx1.^11^ The advantage of using this full E3 over VCB alone in interaction studies with PROTACs is that it is a functionally active version of the E3 that can recruit both the substrate protein (or POI) and the E2-Ub. Specifically, we explore analysis of CRL2^VHL^, the corresponding ternary complexes mediated by PROTACs MZ1 and ACBI3 with their respective POIs BRD4^BD2^ and KRAS,^12^ the CRL2^VHL^-E2-Ub interaction and the full POI-PROTAC-CRL2^VHL^-E2-Ub complexes of both systems. As it is increasingly important to comprehensively characterise such multimeric complexes to more systematically support degrader development^13^ our goal was to capture the interactions of the full E3 with the PROTAC and all the protein partners responsible for ensuring successful Ub transfer to the POI across different systems. We show how this depth of analysis could be achieved rapidly with nMS in a single mass spectrum from a single sample, with simultaneous observation of all equilibrium species. This is not possible with other biophysical methods. We employed *cis-*MZ1 or *cis*-ACBI3 as PROTAC negative controls across the various sample combinations, this further demonstrates the ‘everything is visible’ benefits of nMS analysis for application to targeted protein degradation technologies.

## Results and Discussion

### Native mass spectrometry analysis of individual proteins and binary complexes of first protein of interest-PROTAC system: BRD4^BD2^-MZ1

Native mass spectrometry (nMS) is well developed for characterising biomolecular interactions, including protein-small molecule and protein-protein interactions that underpin larger protein complexes.^14–17^ nMS has been used in a small number of studies to characterise CRLs (without PROTAC)^18, 19^ or PROTAC binary and ternary complexes with a minimal E3,^20–22^ however nMS studies are yet to characterise complexes with a full E3 comprising the Cullin scaffold and RING protein in the context of targeted protein degradation. Given the critical importance of the full E3 species in the mechanism of targeted protein degradation, we wanted to address this gap. Recently, cryo-EM structures of the ternary complex of CRL2^VHL^, MZ1 and BRD4^BD2^ (BRD4^BD2^-MZ1-(NEDD8)-CRL2^VHL^) and the covalently crosslinked (for stability of an otherwise transient species) complex with the E2 UBE2R1 (BRD4^BD2^-MZ1-(NEDD8)-CRL2^VHL^-UBE2R1(C93K)-Ub) were reported (PDB 8RWZ and PDB 8RX0, respectively), **Figure 2**.^23^ When compared to the minimal ternary complex with VCB (PDB 5T35, **Figure 2**) the cryo-EM structures revealed the conserved engagement of BRD4^BD2^ with the VHL component of CRL2^VHL^, in addition to a slightly tilted and reoriented alignment of BRD4^BD2^ distal to the MZ1-VHL binding interface toward Rbx1, reducing the distance between the substrate protein and the catalytic portion of the E3 and the E2-Ub position for ubiquitination.^23^ The observations made from analysis of the cryo-EM structures confirm the significance of using the whole native catalytic enzymatic machinery for investigating more detailed and nuanced mechanistic features for ubiquitination of the PROTAC-recruited target protein. *In vitro* ubiquitination assays and mass spectrometry profiling validated the predominant E2 UBE2R1 ubiquitinated lysine residues of BRD4^BD2^ were consistent with the cryo-EM structures.^23^ The ligand chemical structures of MZ1 and *cis*-MZ1 for this initial nMS study with BRD4^BD2^ are shown in **Figure 2**.

**Figure 2.**
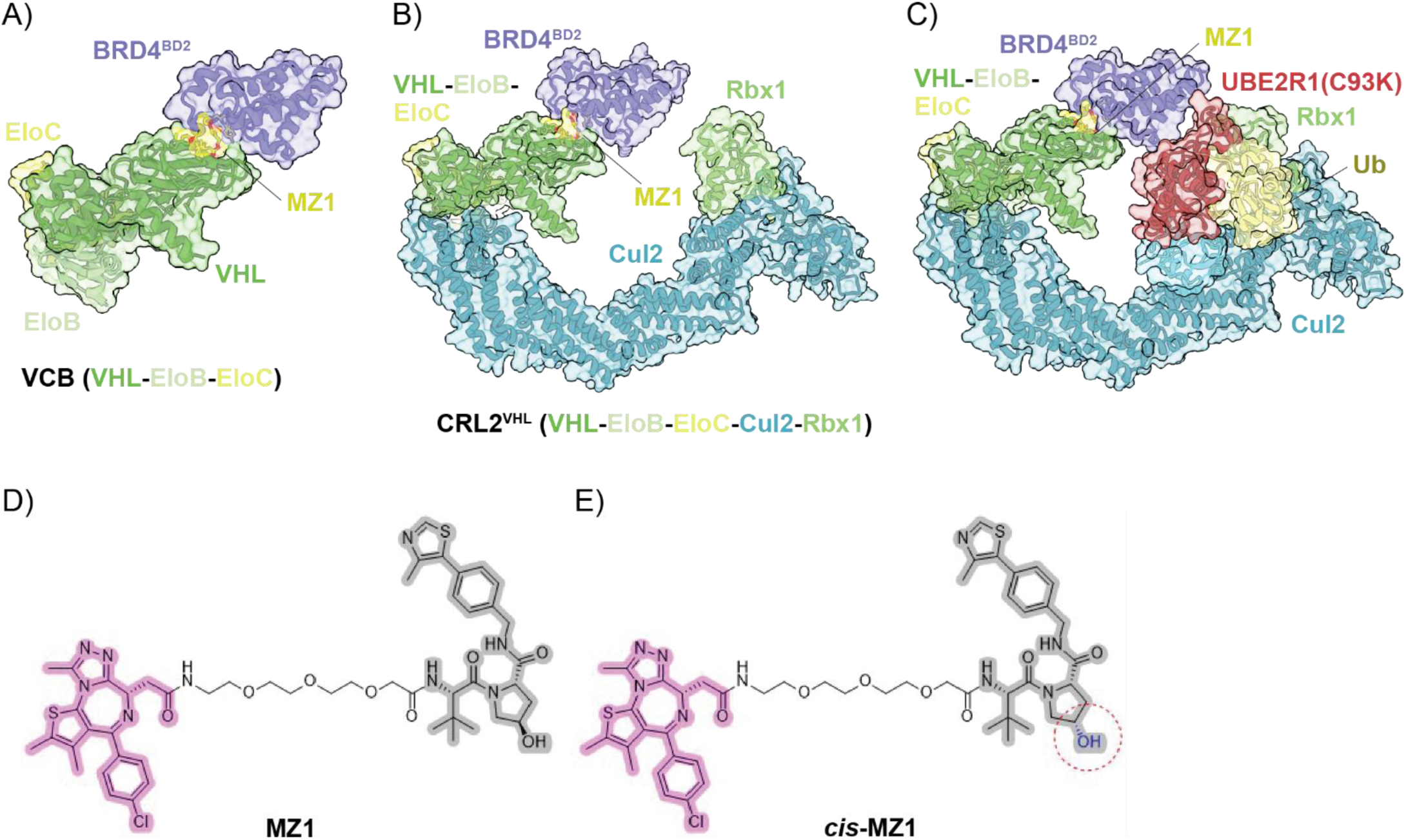
Structures of the protein complexes and PROTAC (with matched negative control) used to establish the feasibility of nMS to characterise all relevant species involved in targeted protein degradation. A) X-ray structure of minimal E3 ternary complex BRD4^BD2^-MZ1-VCB (PDB 5T35). B) Cryo-EM structure of full E3 ternary complex BRD4^BD2^-MZ1-CRL2^VHL^ (PDB 8RWZ). C) Cryo-EM structure of covalently crosslinked BRD4^BD2^-MZ1-CRL2^VHL^-E2-Ub complex (PDB 8RX0).^23, 24^ D) Structures of MZ1, a PROTAC that recruits VHL (grey, pharmacophore from VHL inhibitor VH032) and BRD4^BD2^ (pink, pharmacophore from BRD4^BD2^ inhibitor JQ1), and E) *cis*-MZ1, a negative control with no measurable binding affinity (as determined by ITC)^12^ for VHL owing to the inverted stereochemistry of the hydroxyl group (red circle) compared to MZ1.^12^

Prior to assessment of the sample compositions of interest, the individual protein components BRD4^BD2^, VCB, CRL2^VHL^, UBE2R1(C93K), and UBE2R1(C93K)-Ub (Ub covalently conjugated to the C93K of the E2), were first analysed by nMS to confirm sample identity, assess sample purity and to provide control spectra to support the analysis of the anticipated more complex mass spectra for the mixed samples. All proteins were observed as a single dominant species with high purity, **Supporting Information**, **Figure S1, Tables S1, S2** and **S3**. The measured MW for BRD4^BD2^, VCB and UBE2R1(C93K) agreed with the expected MW based on the protein construct sequences. The measured MW of the pentameric CRL2^VHL^ E3 ligase was ~175.5 Da higher than expected (observed 141,007 Da, expected based on protein construct sequence 140,831.5 Da), this may be attributed to three zinc cations as present in the cryo-EM structures (PDB 8RWZ and PDB 8RX0), or other adducts of this larger multiprotein species. The observed MW of the E2-ubiquitinating conjugate protein with Ub, UBE2R1(C93K)-Ub, was ~964 Da larger than expected (observed 36,247 Da, expected 35,283.5.4 Da) based on the protein sequence and likely corresponded to the addition of extra N-terminal residues of Ub, as previously characterised via denaturing LC-MS.^23^ The LC-MS measured mass was consistent with the MW determined by nMS. The UBE2R1(C93K)-Ub nMS spectrum contained a second minor species (observed 26,630 Da, ~9.6%) that corresponded to the protein without Ub covalently attached (expected MW 26,631 Da), **Supporting Information**, **Figure S1(E)**.

The noncovalent interactions of the protein components comprising VCB (minimal E3) and CRL2^VHL^ (full E3) were assessed with in-source collision-induced dissociation (CID), by increasing the in-source trapping desolvation voltage. Under CID conditions of −75 V, VCB (trimeric VHL-EloC-EloB, 41,373 Da) dissociated into subunits of VHL-EloC (29,639 Da), EloC-EloB (22,696 Da) and VHL (18,675 Da). Free EloC (10,964 Da) and free EloB (11,733 Da) were the only potential dissociation species not observed, **Supporting Information**, **Figure S2** and **Table S4**. Increasing the CID conditions to −150 V resulted in the observation of all possible subunit dissociation products from VCB, these include VHL-EloB (30,408 Da), VHL-EloC (29,639 Da), EloC-EloB (22,695 Da), VHL (18,675 Da), EloB (11,732 Da) and EloC (10,962 Da). In-source CID (−75 V) of CRL2^VHL^ (pentameric VHL-EloB-EloC-Cul2-Rbx1, 141,007 Da) resulted in the observation of Cul2-Rbx (observed 99,583 Da, ~125 Da higher MW than expected MW 99,458 Da), VCB (VHL-EloC-EloB, 41,369 Da), VHL-EloC (29,638 Da), EloC-EloB (22,697 Da), VHL (18,676 Da) and EloB (11,732 Da). An additional 129,257 Da species was also observed that is tentatively assigned as either VHL-EloC-EloB-Cul2 (expected MW 128,558 Da, requires loss of Rbx1 subunit) or VHL-EloC-Cul2-Rbx1 (expected MW 129,098 Da, requires loss of EloB subunit). The latter complex having a MW more consistent with that observed, but an interesting observation if EloB is the subunit lost from this complex as VCB is a stable trimeric species and more investigation is required to assign this species. Free Cul2 (87,184 Da), Rbx1 (12,274 Da) or EloC (10,964 Da, as for VCB dissociation) were not observed, **Supporting Information**, **Figure S2** and **Table S4**. Increasing the CID conditions to −125 V (visibility of protein species was lost at −150 V, thus, not allowing for the direct comparison with VCB), resulted in the observation of the same subunits as described at −75 V as well as the dissociated subunits of Cul2 (89,954 Da), VHL-EloB (30,408 Da), and free EloC (10,962 Da). Additionally, a 113,455 Da species was observed that could not be assigned to any combination of the subunits, therefore, suggesting CID may have induced subunit fragmentation, however further investigation would be required. The only individual protein subunit not observed following the increased desolvation voltage CID from −75 V to −125 V was free Rbx1 (12, 274 Da).

The binary binding of PROTAC MZ1 and the negative control *cis*-MZ1 to BRD4^BD2^ (POI) with the two E3s (VCB and CRL2^VHL^) was next assessed. Both MZ1 and *cis-*MZ1 bound to BRD4^BD2^, **Figure 3** and **Supporting Information**, **Table S5**. This is consistent with the JQ1 pharmacophore present in both compounds (JQ1 is the pan-BET selective bromodomain inhibitor used in development of MZ1^12, 25, 26^), **Figure 2**. In contrast, binding to VCB or CRL2^VHL^ was observed only with MZ1 and not *cis*-MZ1, **Figure 3** and **Supporting Information, Table S5**. This difference is consistent with the inverted stereochemistry of the hydroxyl group of the hydroxyproline moiety of *cis-*MZ1 compared to MZ1 that is known to abolish binding to VHL,^12^ **Figure 2**.

**Figure 3.**
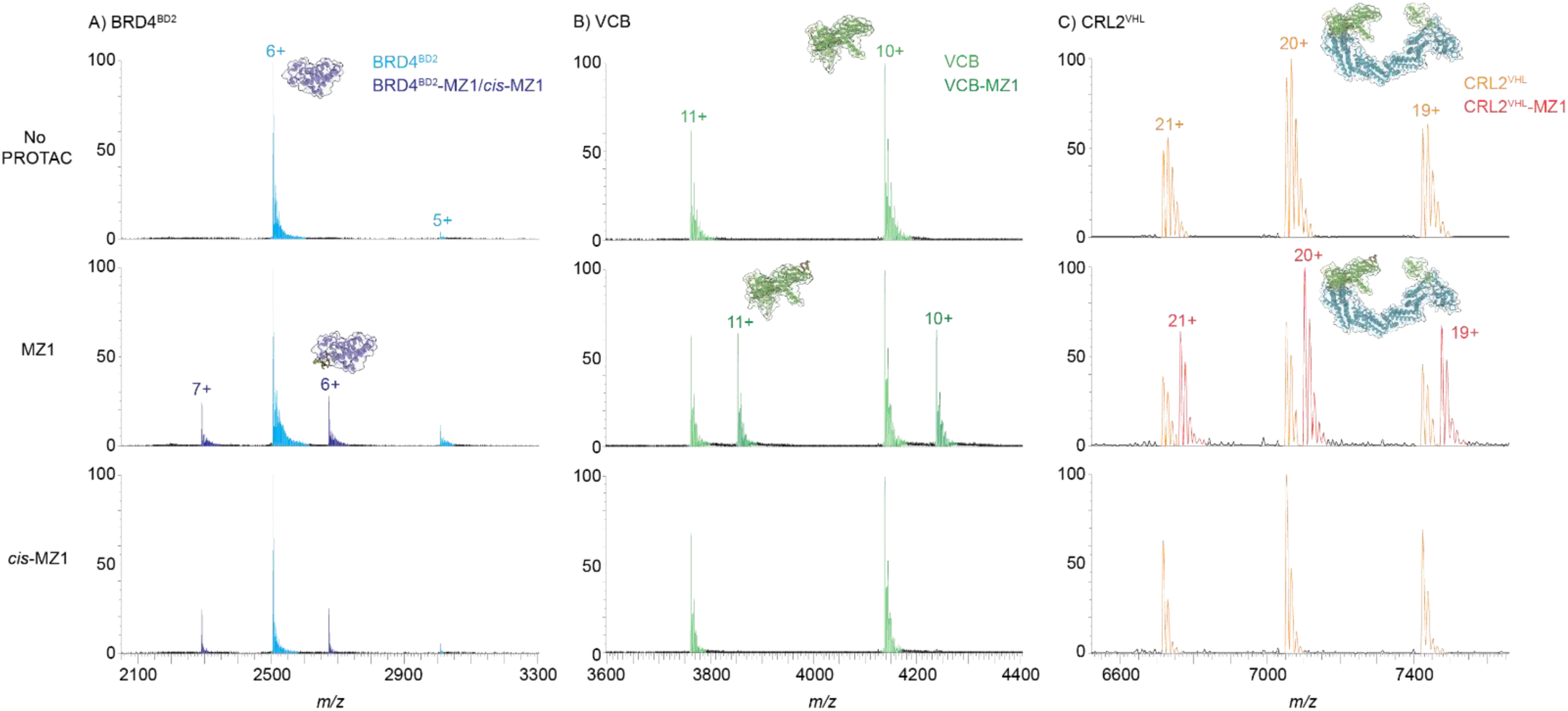
nMS analysis of binary complexes formed by the BRD4^BD2^-MZ1 system. nMS assessment of binary interactions of MZ1 and *cis*-MZ1 (2 µM, 2 equiv) following incubation with 1 µM of A) BRD4^BD2^, B) VCB, or C) CRL2^VHL^ (150 mM NH_4_OAc, 1% DMSO). The expected *m/z* shift corresponding to the binding of PROTAC MZ1 was observed in all samples, while the expected *m/z* shift corresponding to the binding of negative control *cis-*MZ1 was observed only with BRD4^BD2^ and not with either E3. Observed *m/z* values and MWs are in **Supporting Information**, **Table S5**.

### nM analysis of ternary complex formation

For targeted protein degradation, ternary complex formation is critical as the initial target engagement step for subsequent ubiquitination and degradation of the POI. Most methods to screen and characterise potential degrader ligands rely on ternary complex formation as proof of target engagement. Using nMS we observed no interaction of BRD4^BD2^ and VCB without PROTAC, but in the presence of the PROTAC MZ1 the ternary complex BRD4^BD2^-MZ1-VCB was observed (57,409 Da, expected 57,412 Da), consistent with earlier reports,^27, 28^ **Figure 4A** and **Supporting Information**, **Table S6**. The optimal MZ1 concentration for maximising ternary complex formation over the binary interactions with each protein partner (to diminish the hook effect) was 1.4 equiv. This concentration was established by titration of MZ1 (1.0-2.0 equiv, in 0.1 increments) against an equimolar mixture of VCB and BRD4^BD2^, **Supporting Information**, **Figures S3** and **S4.** With 1.4 equiv PROTAC, minimal free VCB or BRD4^BD2^ were observed, with no binary binding to BRD4^BD2^ but reasonable binary binding to VCB (approx. equal relative intensity to unbound VCB) detected, an indicator of a very strong ternary complex. We re-examined this system with *cis*-MZ1 (1.4 equiv) in place of MZ1 and no ternary complex was observed, while unbound and *cis-*MZ1-bound BRD4^BD2^ and unbound VCB were observed, **Figure 4A** and **Supporting Information, Table S6**. This confirms the utility of *cis*-MZ1 as a negative control for MZ1 in nMS analysis.

**Figure 4.**
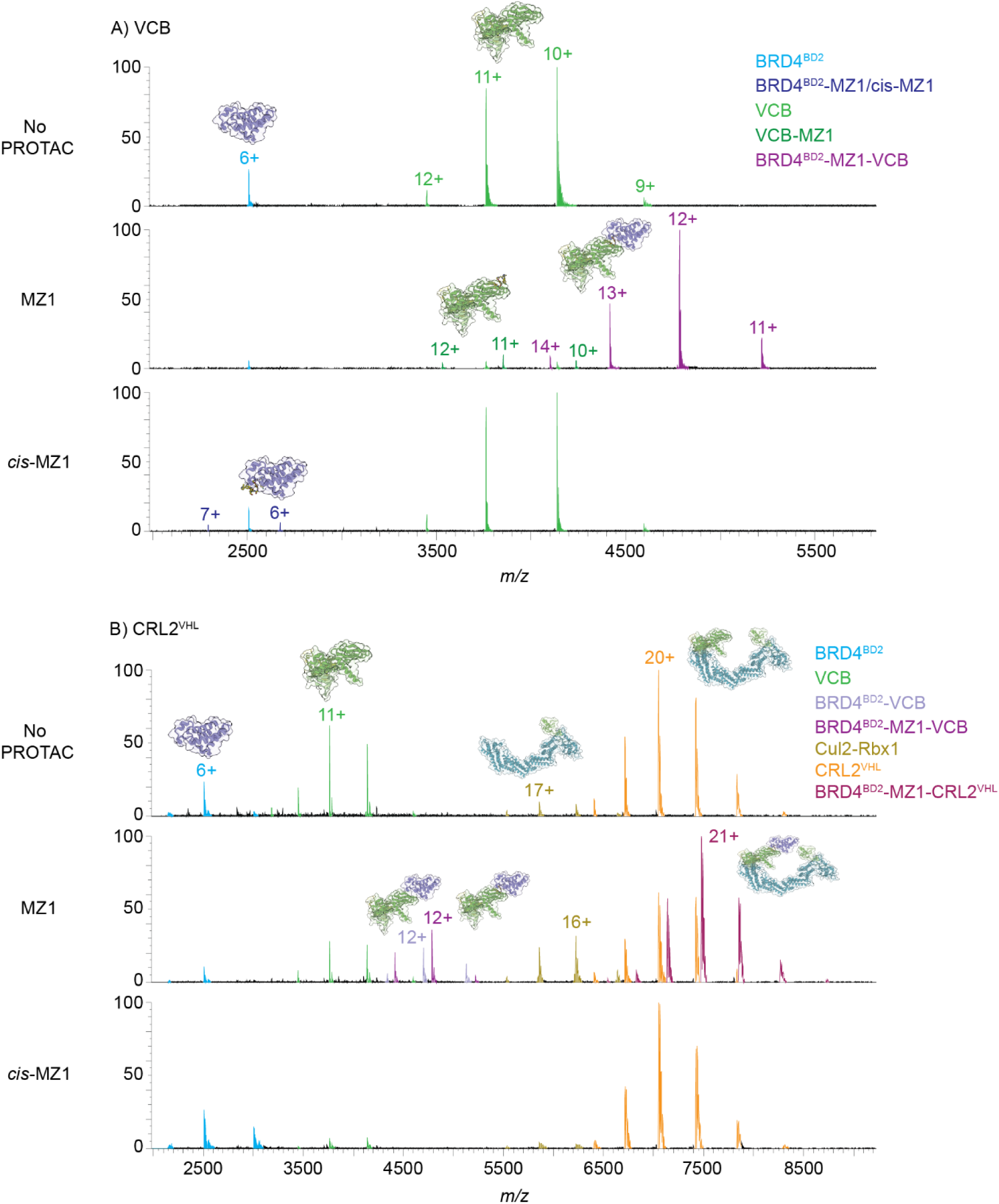
nMS analysis of ternary complexes formed by the BRD4^BD2^-MZ1 system. nMS analysis of ternary complex POI-PROTAC-E3 formation between BRD4^BD2^ (1 µM, 1 equiv), MZ1 (1.4 µM, 1.4 equiv) and A) VCB (1 µM, 1 equiv), or B) CRL2^VHL^ (1 µM, 1 equiv) (150 mM NH_4_OAc, 1% DMSO). Mass spectra are raw data with charge state (all for VCB and most intense only for each species for CRL2^VHL^) and species identity annotated. Top panel is no PROTAC control (1% DMSO), middle panel is with addition of MZ1, and lower panel is with addition of negative control *cis*-MZ1 (1.4 equiv). Observed *m/z* values and MWs are in **Supporting Information**, **Table S6**.

We then assessed if the ternary complex with the full E3 CRL2^VHL^ was observable by nMS. In the absence of PROTAC, there was no interaction between CRL2^VHL^ and BRD4^BD2^, similarly to the corresponding minimal E3 VCB sample, **Figure 4B** and **Supporting Information, Table S6**. Both free BRD4^BD2^ and CRL2^VHL^ were observed, in addition to minor quantities of VHL-EloC-EloB and Cul2-Rbx1, indicating some dissociation of CRL2^VHL^. On addition of MZ1 (1.4 equiv), the ternary complex comprising BRD4^BD2^-MZ1-CRL2^VHL^ was the major species observed (157,099 Da, expected 157,046 Da), **Figure 4B** and **Supporting Information, Table S6**. There was some free CRL2^VHL^ observed indicating not all the intact E3 had been recruited into the ternary complex, while lesser amounts of the dissociated subunits VHL-EloC-EloB and Cul2-Rbx1 were also present. Interestingly, there was some ternary complex where the CRL2^VHL^ had dissociated to give VCB (VHL-EloC-EloB) but the ternary complex remained intact (i.e. BRD4^BD2^-MZ1-VCB). Further to this, the complex BRD4^BD2^-VCB was also observed wherein the PROTAC had been dissociated, a phenomenon previously seen in the gas phase nMS measurement of PROTAC ternary complexes.^28–31^ When replacing MZ1 with the negative control *cis*-MZ1, no ternary complex was observed, however, unbound BRD4^BD2^, unbound CRL2^VHL^ and unbound dissociated VCB and Cul2-Rbx1 were observed. *cis-*MZ1-bound BRD4^BD2^ was not observed, however it may be too low in intensity relative to background noise in these spectra, **Figure 4B** and **Supporting Information, Table S6**.

### Assessment of the E3-E2 interaction with nMS

The CRL class of E3s transiently recruit an E2 linked to Ub to facilitate Ub transfer to a protein substrate for degradation.^1, 8^ nMS analysis of a mixture of CRL2^VHL^ (1 equiv) and an E2 covalently preloaded with Ub, UBE2R1(C93K)-Ub (5 equiv, the Cys to Lys mutation enables covalent loading of a donor Ub^23^) revealed the formation of an E3-E2-Ub complex between CRL2^VHL^ and UBE2R1(C93K)-Ub (177,339 Da, expected 177,254 Da), **Figure 5A, Supporting Information Figure S5** and **Table S7**. The peak intensity for the complex was significant (~15.3% relative intensity), however the major species observed were the free CRL2^VHL^ (with some dissociated VCB) and free E2 UBE2R1(C93K)-Ub (and the minor UBE2R1(C93K) component without Ub conjugated), **Figure 5A**. In a physiological setting, once E2 transfers the Ub cargo to the substrate, it must dissociate from the E3 to be replenished with the next E2-Ub, hence this interaction is transient by nature^8^ and therefore exciting to see this interaction was observable by nMS, noting it was not possible to capture this interaction by cryo-EM without covalent crosslinking to form the BRD4^BD2^-MZ1-CRL2^VHL^-Ub-E2 conjugated species to artificially increase the occupancy of the E2 on the E3 (PDB 8RXO).^23^

**Figure 5.**
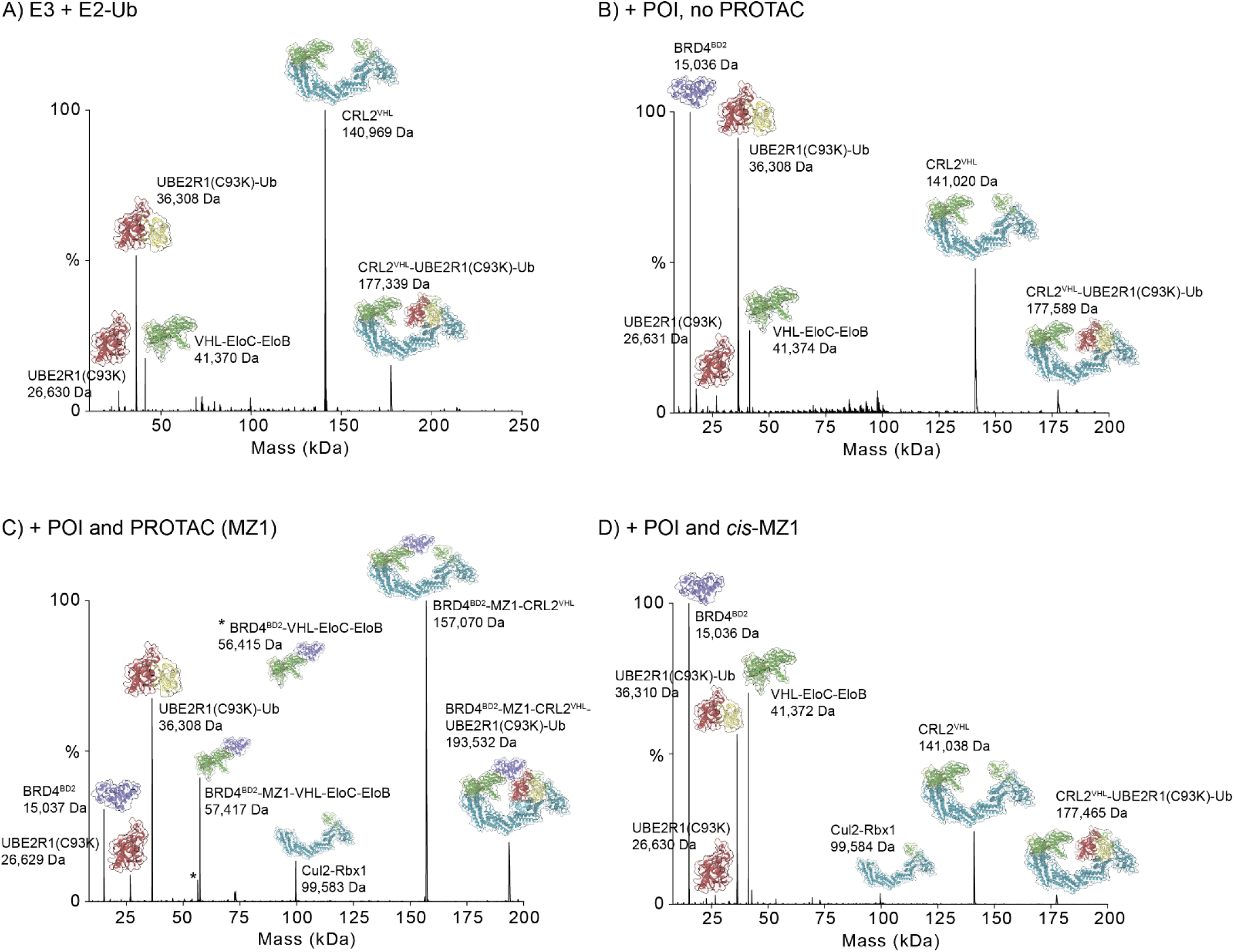
nMS assessment of formation of whole complex and components of BRD4^BD2^-MZ1-CRL2^VHL^-Ub-E2-Ub system. A) Deconvoluted nMS spectrum of CRL2^VHL^ (1 µM, 1 equiv) in the presence of UBE2R1(C93K)-Ub (5 µM, 5 equiv, 150 mM NH_4_OAc). The transient E3-E2-Ub interaction CRL2^VHL^-UBE2R1(C93K)-Ub was observable (177,339 Da). Free CRL2^VHL^ was the major peak observed, with dissociated VHL-EloC-EloB and UBE2R1(C93K)-Ub also observed (with minor UBE2R1(C93K) contaminant). The corresponding raw data mass spectra and observed *m/z* values and MWs are shown in **Supporting Information**, **Figure S5** and **Table S7**, respectively. B-D) Deconvoluted nMS spectra showing PROTAC mediated formation of the BRD4^BD2^-MZ1-CRL2^VHL^-UBE2R1(C93K)-Ub complex. B) No PROTAC control (0.69% DMSO), C) with MZ1 (1.5 equiv, 1.39 µM), D) with *cis*-MZ1 (1.5 equiv, 1.39 µM). Protein component ratios of 1.5 equiv BRD4^BD2^ (1.39 µM), 1.5 equiv PROTAC/negative control (1.39 µM), 1 equiv CRL2^VHL^ (0.92 µM), 5 equiv UBE2R1(C93K)-Ub (4.63 µM) (with final 0.69% DMSO). Raw data mass spectra and observed *m/z* values and MWs are shown in **Supporting Information**, **Figure S6** and **Table S8**.

The formation of an E3-E2 complex without Ub preloaded onto the E2 was also assessed. Here, UBE2R1(C93K) (2 equiv) was added to CRL2^VHL^ (1 equiv) and a minor amount of E3-E2 complex was observed (167,632 Da, expected 167,638 Da, 5.9% relative intensity), this was a similar intensity to the complex formed when UBE2R1(C93K)-Ub (2 equiv) was added (3.9% relative intensity) (data not shown). This indicated the transient E3-E2 interactions in this system can be observed regardless of the loading with Ub, despite previous observations that the E3-E2 interaction is stronger in the presence of Ub.^32^

### Assessment of the full BRD4^BD2^-MZ1-CRL2^VHL^-UBE2R1(C93K)-Ub complex with nMS

Next, we investigated the feasibility of capturing the full degradation complex induced by the PROTAC using nMS. For this analysis the sample components were combined in ratios to replicate the conditions used for the cryo-EM study of the related system, **Figure 2**.^23^ nMS analysis of the mixture of BRD4^BD2^ (1.5 equiv), the full E3 CRL2^VHL^ (1 equiv) and the Ub-conjugated E2 UBE2R1(C93K)-Ub (5 equiv) in the absence of PROTAC (0.69% DMSO) showed a low intensity peak corresponding to the CRL2^VHL^-UBE2R1(C93K)-Ub (i.e. E3-E2-Ub) complex (observed 177,589 Da, expected 177,465 Da, ~7.7 %, relative intensity), **Figure 5B, Supporting Information, Figure S6** and **Table S8**. This was consistent with that observed when only the E3 and E2 were combined as described in the previous section, **Figure 5A**. Free BRD4^BD2^, CRL2^VHL^ and UBE2R1(C93K)-Ub were readily detected, along with some dissociated VCB from the full E3 CRL2^VHL^, **Figure 5B**. Addition of PROTAC MZ1 (1.5 equiv) to this sample resulted in the observation of BRD4^BD2^-MZ1-CRL2^VHL^-UBE2R1(C93K)-Ub complex (i.e. POI-PROTAC-E3-E2-Ub) (193,479 Da, expected 193,293 Da, 19.6% relative intensity), **Figure 5C**. This corresponds to the full PROTAC-engaged degradation complex that is essential for transferring Ub from the E2 onto the POI. Other species detected in the native mass spectrum included the PROTAC-mediated ternary complex BRD4^BD2^-MZ1-CRL2^VHL^ and various dissociated ternary complex species including BRD4^BD2^-MZ1-VHL-EloC-EloB and this complex with the PROTAC dissociated. Free BRD4^BD2^, free E2-Ub UBE2R1(C93K)-Ub (as well as E2 UBE2R1(C93K) without Ub loaded) are observed. Interestingly no free E3 CRL2^VHL^ was observed, however, the dissociated subunit complex Cul2-Rbx1 was observed. MZ1 was then replaced with *cis*-MZ1, and the results essentially mimicked the DMSO only control, **Figure 5D**. A minor amount of E3-E2-Ub (CRL2^VHL^-UBE2R1(C93K)-Ub) was detected (177,465 Da, ~3.2% relative intensity), however, no ternary complex or ternary complex engaged with E2-Ub was observed. This is consistent with the structural difference of *cis-*MZ1 compared to MZ1 that abolishes VHL-substrate protein binding.^12^ The only protein species observed by nMS from this sample were the individual free components BRD4^BD2^, CRL2^VHL^ and UBE2R1(C93K)-Ub (and contaminating UBE2R1(C93K)), as well as some dissociated VCB and Cul2-Rbx1 from the full E3 CRL2^VHL^.

### Assessment of key interactions of a second protein of interest-PROTAC system KRAS-ACBI3 with full E3 ligase CRL2^VHL^ using nMS

To assess the scope of nMS beyond the BRD4^BD2^-MZ1 POI-PROTAC system described above, we investigated the interactions of a second example POI-PROTAC system, specifically wild type KRAS protein and the pan-KRAS PROTAC ACBI3, which leads to potent degradation of many KRAS variants mediated by VHL.^33^ There is a cryo-EM structure of the KRAS-ACBI3-CRL2^VHL^ complex (PDB 8QU8), however, there are no structures of the higher level complexes with E2-Ub reported, **Figure 6A**.^33^ Similar to MZ1, ACBI3 comprises the VHL recruiting ligand, while the matched negative control compound *cis*-ACBI3, a stereoisomer that binds KRAS but is unable to bind VHL, was used in our nMS analysis, **Figure 6B, 6C**. nMS analysis of KRAS confirmed the protein was a single dominant species with an observed MW of 19,882 Da, this is ~32 Da higher than the expected MW based on the GDP-bound form of the protein sequence (19,850.4 Da), indicative of a possible methionine oxidation, **Supporting Information, Figure S7**. The binary binding of PROTAC ACBI3 and negative control *cis*-ACBI3 to KRAS (POI) and the two E3 constructs, VCB and CRL2^VHL^, was assessed. As expected, both ACBI3 and *cis*-ACBI3 bound to KRAS, while only ACBI3, and not *cis*-ACBI3, bound to VCB and CRL2^VHL^, **Supporting Information, Figure S7** and **Table S9**. A second smaller species (MW ~515 Da) was observed binding to VCB and CRL2^VHL^ in the samples where ACBI3 was added. An independent LC-MS analysis of the ACBI3 compound stock obtained from opnMe confirmed the presence of this impurity (data not shown) that may result from chemical synthesis or compound degradation on storage.

**Figure 6.**
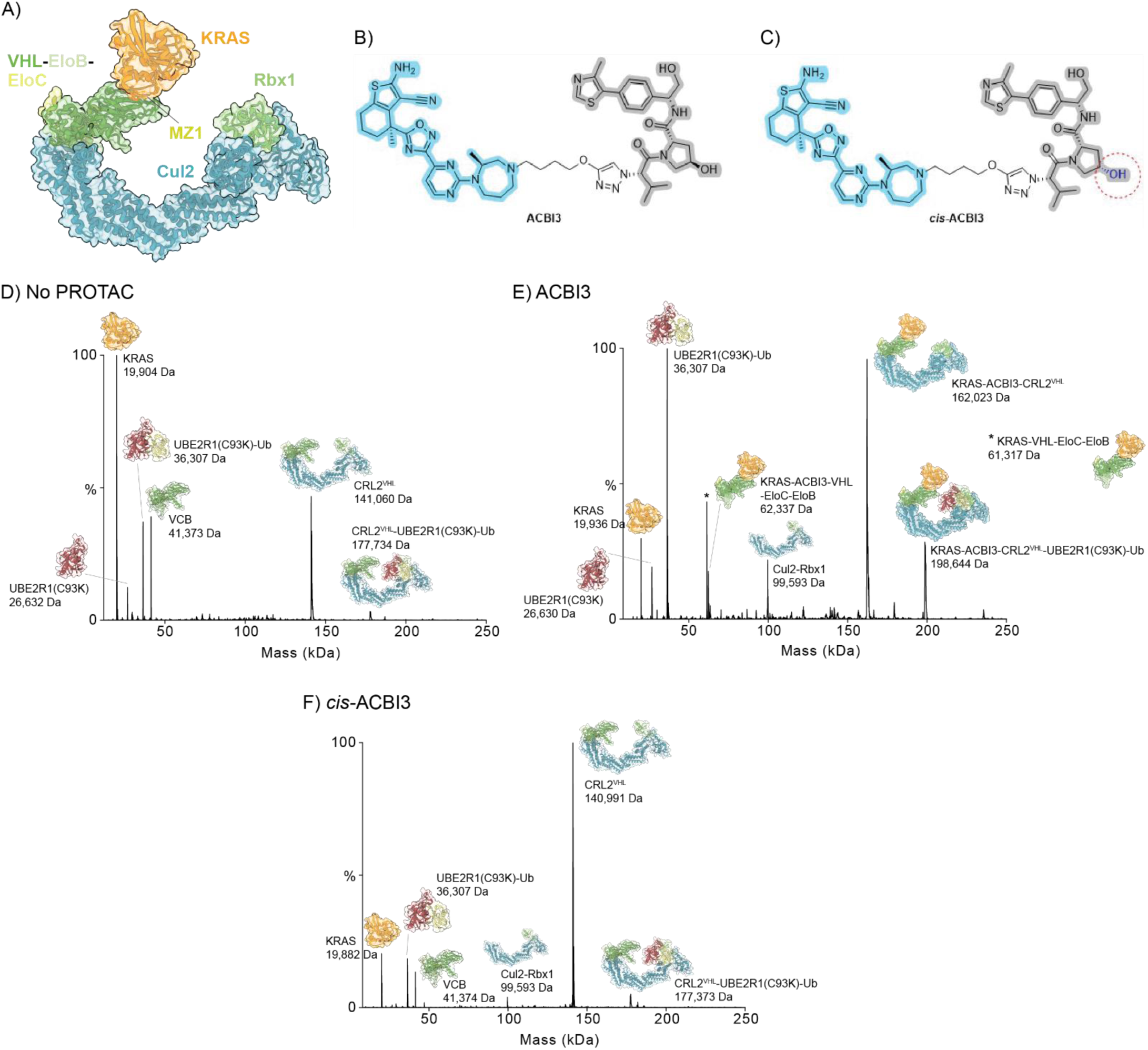
nMS assessment of formation of whole complex and components of KRAS-ACBI3-CRL2^VHL^-Ub-E2-Ub system. A) X-ray structure of full E3 ternary complex KRAS-ACBI3-CRL2^VHL^ (PDB 8QU8).^33^ B) Chemical structure of ACBI3, a PROTAC that recruits VHL (grey, pharmacophore from VHL inhibitor VH032) and KRAS (blue pharmacophore present in KRAS inhibitors). C) Chemical structure of matched negative control ligand *cis-*ACBI3 with inverted stereochemistry of the hydroxyl group (red circle) and no measurable binding for VHL but retained binding to KRAS.^33^ D-F) Deconvoluted nMS spectra showing PROTAC mediated formation of the KRAS-ACBI3-CRL2^VHL^-UBE2R1(C93K)-Ub complex. D) no PROTAC control (0.75% DMSO), E) with ACBI3 (1.9 equiv, 1.9 µM), F) with *cis*-ACBI3 (1.9 equiv, 1.9 µM). Protein component ratios of 1 equiv KRAS (1 µM), 1.9 equiv PROTAC/negative control (1.9 µM), 1 equiv CRL2^VHL^ (1 µM), 5 equiv UBE2R1(C93K)-Ub (5 µM) (with final 0.75% DMSO). Raw data of mass spectra and observed *m/z* values and MWs are shown in **Supporting Information**, **Figure S11** and **Table S12**, respectively.

nMS analysis of KRAS (1 equiv) and VCB (1 equiv) or CRL2^VHL^ (1 equiv) showed no interaction between the two proteins, with only unbound proteins observed and, for the full E3, also some VCB and Cul2-Rbx1 dissociation products, **Supporting Information, Figure S8** and **Table S10** and **S11**. A titration of ACBI3 (0.8-2.5 equiv, 0.1 increments) against the equimolar mixture of KRAS and VCB (1 µM each) revealed the optimal ratio of PROTAC for ternary complex observation was 1.9 equiv, **Supporting Information Figure S9** and **S10**. Of note, significant binary binding of PROTAC to VCB was also observed concurrent with ternary complex formation, this differs to the order of complex formation for MZ1 where strong ternary complex formation was observed prior to significant VCB binary binding. The presence of the impurity in the ACBI3 sample would influence the ACBI3 stock concentration and could alter the ternary complex formation dynamics. On addition of ACBI3 (1.9 equiv) the KRAS-ACBI3-VCB ternary complex forms as the most intense species observed in the mass spectrum, whilst the KRAS-ACBI3-CRL2^VHL^ ternary complex formed with similar relative intensity to the free CRL2^VHL^, **Supporting Information, Figure S8** and **Table S11**. No ternary complex formation was observed when *cis-*ACBI3 was added, with mass spectra similar to those with no PROTAC added, **Supporting Information, Figure S8** and **Table S11**.

The full POI-PROTAC-E3-E2-Ub ubiquitination complex was then assessed, **Figure 6D-F** and **Supporting Information, Figure S11** and **Table S12**. In the absence of ACBI3, a minor amount of E3-E2-Ub was observed (CRL2^VHL^-UBE2R1(C93K)-Ub, relative intensity 3.3%) together with free KRAS, UBE2R1(C93K)-Ub (and contaminating UBE2R1(C93K)), and CRL2^VHL^ (including dissociation products VCB and VHL-EloC), **Figure 6D**. On addition of ACBI3 (1.9 equiv), the KRAS-ACBI3-CRL2^VHL^-UBE2R1(C93K)-Ub complex was detected (relative intensity 28.4%), as well as ternary complexes KRAS-ACBI3-CRL2^VHL^ and KRAS-ACBI3-VCB (where VCB had dissociated from the CRL2^VHL^ but the ternary complex interactions remained) and KRAS-VCB (from dissociation of the PROTAC) were observed, **Figure 6E**. Additionally, free KRAS, UBE2R1(C93K)-Ub (and contaminating UBE2R1(C93K)) and dissociated Cul2-Rbx1 were also detected. When *cis*-ACBI3 (1.9 equiv) was added instead of ACBI3, a minor amount of the E3-E2-Ub complex was observed (CRL2^VHL^-UBE2R1(C93K)-Ub, relative intensity 5.1%), with free KRAS, UBE2R1(C93K)-Ub and CRL2^VHL^ (and dissociated VCB and Cul2-Rbx1) also detected, as for the DMSO control, **Figure 6F**. Overall, the behaviour of this second POI system with PROTAC and matched negative control was similar to the BRD4 system with MZ1 and cis-MZ1, indicative of the scope and applicability of the nMS method to different degrader systems.

## Conclusions

Many degraders in development act through the pentameric Cullin RING E3 ligase complex CRL2^VHL^ yet there are few biophysical studies that have characterised this species, with most focusing on the nonfunctional minimal E3 VCB without the Cullin scaffold or RING proteins. Herein we demonstrate the characterisation of CRL2^VHL^ and all expected full E3 complexes integral to the targeted protein degradation mechanism using nMS. This includes the binary, ternary and higher order multiprotein complexes made up of the full E3 CRL2^VHL^, POI (BRD4^BD2^ or KRAS) and E2-Ub with and without addition of the PROTAC (MZ1 or ACBI3) and covers highly stable complexes through to those expected to be transient and/or weak complexes. We establish direct insight into the stoichiometry of interactions and observe all equilibrium species within a sample in the same measurement, including the expected species and species not readily predicted to form. These unprecedented advantages of nMS set it apart from other biophysical methods where this is not possible. Whilst high-resolution complex structures provide crucial mechanistic insights into degrader-driven interactions that underpin successful target ubiquitination, the employment of X-ray crystallography or cryo-EM it is not always possible and is not routine. These methods commonly require troubleshooting, often against modified protein constructs (including covalent crosslinking to stabilise complexes) to support data collection. As a result, so far there are only a handful of structures of a PROTAC ternary complex wherein a full E3 is used, let alone structures of the higher order interactions with E2 and Ub, despite the intensity of research in the field.^23, 33, 34^ Given the depth of information achieved from nMS analysis that is without resource intensive sample preparation requirements other than standard purified native proteins and was readily applicable across two different systems, we suggest a greater role for nMS in the assessment of multiprotein complex formation to triage degraders, POIs and/or full E3s that are in development for further structural analysis by X-ray crystallography or cryo-EM and/or functional cell-based investigation, with the overall impact of accelerating targeted protein degradation technologies. Future nMS studies could add further key protein components to the analysis, including NEDD8 modification of the CRL, a protein component important for activation of the CRL to facilitate Ub transfer.^35^

## Methods

### Protein and ligand samples

BRD4^BD2^, VCB, UBE2R1(C93K) and UBE2R1(C93K)-Ub samples were obtained from a previous study.^23^ CRL2^VHL^ and KRAS (GDP-bound form) protein production is provided in **Supporting Information**. The test compounds MZ1 and ACBI3 (PROTACs) and the matched negative controls *cis*-MZ1 and *cis-*ACBI3 were provided as dry stocks from opnMe (https://www.opnme.com/), the open science portal of Boehringer Ingelheim.^36^ The compounds were solubilised to 10 mM in DMSO and the compound stock solution stored at −80 °C.

### Native mass spectrometry sample preparation

Proteins were individually buffer exchanged into 150 mM ammonium acetate pH ~6.7 (Thermo Fisher AM9070G) using Micro Bio-Spin P-6 columns (Bio-Rad) according to manufacturer’s protocols. Protein concentration following buffer exchange was determined by measuring the absorption at 280 nm ((NanoDrop; Thermo Fisher) and considered the protein’s molecular weight and extinction coefficient (protein construct sequences provided in **Supporting Information**, **Table S1**). Compound stocks (10% DMSO) were prepared from the parent 100% DMSO stocks to yield a concentration that allowed a further 1/10 dilution into the nMS sample to give the desired compound test concentration for nMS at 1% DMSO (or less). Proteins were diluted in 150 mM ammonium acetate pH ~6.7 to the required concentrations and mixed with their protein binding partners and compounds as required, dependent on the test sample composition. Final concentrations of all test components are indicated for each sample/spectrum. All samples were incubated for at least 30 minutes at room temperature prior to nMS analysis.

### Native mass spectrometry acquisition and data analysis

All nMS analyses was completed using a Q-Exactive Ultra High Mass Range Hybrid Quadrupole-Orbitrap Mass Spectrometer (UHMR MS, Thermo Fisher Scientific). Samples were introduced in the positive ion mode via either nanoelectrospray ionisation (nanoESI) from commercial borosilicate emitters with Au/Pd double coating (Thermo Fisher Scientific ES388, 1.1 mm tip length) or using a TriVersa NanoMate nanoESI source (Advion Biosciences) with a chip containing 5 µm diameter nozzles. nanoESI parameters for the borosilicate emitters included a spray voltage of 0.8-1.4 kV, and for the NanoMate source included a voltage of 1.7 kV and gas pressure of 1 psi. UHMR MS parameters were optimised to maximise signal intensity and signal-to-noise whilst maintaining intact folded proteins and protein complexes, and were in the range of values as follows: resolution 6,250 to 200,000, scan ranges 1,500-6,500, 1,500-8,000 or 2,000-10,000 *m/z*, microscans 5 or 10, averaging 5 or 20, capillary temperature 230 to 275 °C, S-lens RF level 200, desolvation voltage 0 to −60 V, trapping gas pressure 1.0 to 4.0, extended trapping 1.0, ion transfer target *m/z* set for high *m/z*, detector *m/z* optimisation set for low *m/z*, nitrogen as the collision gas, data acquired for at least 1-2 minutes. Specific parameters for methods used for measuring different proteins and/or protein complexes are detailed in **Supporting Information**, **Table S13**. nMS data were analysed directly using Themo Fisher Scientific FreeStyle 1.8 SP2 QF1 and deconvoluted in UniDec.^37^ UniDec data processing parameters were as follows: *m/z* range set to the full *m/z* range acquired, 3 or 4 to 7-40 charge state range, 5,000 or 10,000 to 50,000-200,000 Da MW range (charge state and MW range dependent on the sample), sample mass every 1 Da, peak detection range and threshold of 500 Da and 0.05 or 0.1, respectively. For accurate MW determination, the peak detection range was iteratively reduced to as low as 10 Da to identify the MW of the left-most unadducted peaks. To further identify low intensity species in deconvoluted mass spectra, the peak threshold was lowered to 0.02-0.03 as required. Specific UniDec deconvolution parameters for each protein and/or protein complexes measured are detailed in **Supporting Information**, **Table S13**.

## Safety Statement

No unexpected or unusually high safety hazards were encountered.

## Supporting Information

Supporting material includes: recombinant CRL2^VHL^ and KRAS protein production methods; UHMR MS parameters; protein construct sequences, calculated MWs and corrected expected MWs for each protein and protein complex; supporting nMS raw and deconvoluted mass spectra; observed *m/z* values, charge states and MWs for raw and deconvoluted spectra.

## Data Availability

The data supporting this study are available in this article and the accompanying online supplementary material. Any addition data is available upon reasonable request from the corresponding authors.

## Conflicts of Interest

A.C. is a scientific founder and shareholder of Amphista Therapeutics, a company that is developing targeted protein degradation therapeutic platforms. A.C. is on the Scientific Advisory Board of ProtOS Bio and TRIMTECH Therapeutics. The Ciulli laboratory receives or has received sponsored research support from Almirall, Amgen, Amphista Therapeutics, Boehringer Ingelheim, Eisai, Merck KaaG, Nurix Therapeutics, Ono Pharmaceutical and Tocris Bio-Techne. All other authors declare they have no competing interests.

## Funding

The authors greatly acknowledge the support of the Clive and Vera Ramaciotti Foundation (Biomedical Research Award, grant number 2023BRA19) and the Australian Government through an Australian Research Council’s Industrial Transformation Training Centre grant (grant number IC180100021). Research in the Ciulli laboratory on Cullin RING ligases and targeted protein degraders, including PROTACs, has received funding from the European Research Council (ERC) under the European Union’s Seventh Framework Programme (FP7/2007–2013) as a Starting Grant to A.C. (grant agreement no. ERC-2012-StG-311460 DrugE3CRLs), and from the Innovative Medicines Initiative 2 (IMI2) Joint Undertaking under Grant 875510 (EUbOPEN project). The IMI2 Joint Undertaking receives support from the European Union’s Horizon 2020 research and innovation program, EFPIA companies, and associated partners: KTH, OICR, Diamond, and McGill. C. Crowe was supported by a PhD studentship from the UK Medical Research Council (MRC) under the Industrial Cooperative Awards in Science & Technology (iCASE award with Tocris Bio-Techne) doctoral training programme MR/R015791/1 and is now a postdoctoral researcher funded by the Michael J. Fox Foundation. L. Wieske received funding from the Swedish Research Council (2024-00197) and the IF:Stiftelse, a Swedish foundation for pharmaceutical research.

## Acknowledgements

MZ1, *cis-*MZI, ACB13 and *cis*-ACB13 were kindly provided by Boehringer Ingelheim via its open innovation platform opnMe, available at https://www.opnme.com. We thank Dr Wendy Loa for mass spectrometry instrumentation support and acknowledge the Ramaciotti Australian Native Mass Spectrometry Platform for Health Discoveries, Griffith University. We thank technical support staff at the University of Dundee for the upkeep of protein production facilities. We thank Dr Kevin Haubrich and Dr Yuting Cao (Ciulli lab) for the gift of the KRAS expression plasmid. Molecular graphics of proteins were generated with UCSF ChimeraX, developed by the Resource for Biocomputing, Visualization, and Informatics at the University of California, San Francisco, with support from National Institutes of Health R01-GM129325 and the Office of Cyber Infrastructure and Computational Biology, National Institute of Allergy and Infectious Diseases.^38^

## Notes

### Competing Interest Statement

Alessio Ciulli is a scientific founder and shareholder of Amphista Therapeutics, a company that is developing targeted protein degradation therapeutic platforms. A.C. is on the Scientific Advisory Board of ProtOS Bio and TRIMTECH Therapeutics. The Ciulli laboratory receives or has received sponsored research support from Almirall, Amgen, Amphista Therapeutics, Boehringer Ingelheim, Eisai, Merck KaaG, Nurix Therapeutics, Ono Pharmaceutical and Tocris Bio-Techne. All other authors declare they have no competing interests.

### Summary of Updates

The manuscript revision includes characterisation of a second PROTAC system using native mass spectrometry: There are now two protein of interest examples (BRD4BD2 and KRAS) with their respective PROTACs (MZ1 and ACBI3) and negative controls (cis-MZ1 and cis-ACBI3).

