## Supplementary Information for "Native Mass Spectrometry Analysis of Cullin RING Ubiquitin E3 Ligase Complexes in the Context of Targeted Protein Degradation"

### Table of Contents

**Supporting Table S1.** Sequences and calculated molecular weight (MW) of protein constructs utilised in this study. ....................S4

**Supporting Figure S1.** Raw and deconvoluted nMS spectra for individual protein components. ....................S7

**Supporting Table S2.** Observed *m/z* values, charge states and measured MWs for individual protein components. ....................S8

**Supporting Table S3.** Reference corrected MW of protein components, PROTAC/compound MWs and expected MW of protein complexes determined (based on corrected MW). ....................S9

**Supporting Figure S2.** Collision-induced dissociation (CID) of minimal E3 (VCB) and full E3 ($CRL2^{VHL}$) into individual component subunits. ....................S10

**Supporting Table S4.** Observed MW and assignment to subunits dissociated from E3s VCB and $CRL2^{VHL}$ with in-source CID. ....................S11

**Supporting Table S5.** Observed *m/z* values and measured MWs for binary binding of MZ1 (MW 1002.6 Da) and *cis*-MZ1 (MW 1002.6 Da) to BRD4$^{BD2}$, VCB and CRL2$^{VHL}$. ....................................................S12

**Supporting Table S6.** Observed *m/z* values and measured MWs for samples comprising BRD4$^{BD2}$ and either VCB or CRL2$^{VHL}$ with MZ1 (MW 1002.6 Da) or *cis*-MZ1 (MW 1002.6 Da). ...............................S14

**Supporting Figure S3.** nMS titration of PROTAC MZ1 (1.0-2.0 equiv, 0.1 increments) against POI BRD4$^{BD2}$ and E3 VCB (present at equimolar concentrations, 1 μM each) to determine the optimal MZ1 equiv for maximal ternary complex formation. ...........................................................................................S17

**Supporting Figure S4.** Quantification of ternary complex formation between BRD4$^{BD2}$ and VCB (present at equimolar concentrations, 1 μM) with MZ1 titrated (1.0-2.0 equiv, 0.1 increments) as measured by nMS. .......................................................................................................................................................S18

**Supporting Figure S5.** Raw data nMS spectra of E3-E2-Ub sample comprising CRL2$^{VHL}$ (1 μM) and UBE2R1(C93K)-Ub (5 μM). ...............................................................................................................S19

**Supporting Table S7.** Observed *m/z* values, charge states and measured MWs for interacting species in the nMS spectra of the E3-E2-Ub sample (CRL2$^{VHL}$ and UBE2R1(C93K)-Ub). ............................................S20

**Supporting Figure S6.** Raw data nMS spectra showing PROTAC mediated formation of the full BRD4$^{BD2}$-MZ1-CRL2$^{VHL}$-UBE2R1(C93K)-Ub complex. ...........................................................................................S21

**Supporting Table S8.** Observed *m/z* values and measured MWs for the full POI-PROTAC-E3-E2-Ub degradation complex. .............................................................................................................................S22

**Supporting Figure S7.** nMS assessment of KRAS and binary binding with ACBI3 and *cis*-ACBI3. .......S24

**Supporting Table S9.** Observed *m/z* values and measured MWs for binary binding of ACBI3 (MW 1019.3 Da) and *cis*-ACBI3 (MW 1019.3 Da) to KRAS, VCB and CRL2$^{VHL}$. ....................................................S25

**Supporting Table S10.** Expected MW of KRAS protein complexes (based on corrected MW). ...............S27

**Supporting Figure S8.** nMS analysis of ternary complex POI-PROTAC-E3 formation between KRAS (1 μM, 1 equiv), ACBI3 (1.9 μM, 1.9 equiv) and A) VCB (1 μM, 1 equiv) or B (1 μM, 1 equiv). ................S28

**Supporting Table S11.** Observed *m/z* values and measured MWs for samples comprising KRAS and either VCB or CRL2$^{VHL}$ with ACBI3 (MW 1019.3 Da) and *cis*-ACBI3 (MW 1019.3 Da). .................................S29

**Supporting Figure S9.** nMS titration of PROTAC ACBI3 (0.8-2.5 equiv, 0.1 increments) against POI KRAS and E3 VCB (present at equimolar concentrations, 1 μM each) to determine the optimal ACBI3 equiv for maximal ternary complex formation. ...........................................................................................S31

**Supporting Figure S10.** Quantification of ternary complex formation between KRAS and VCB (present at equimolar concentrations, 1 μM each) with ACBI3 titrated (0.8-2.5 equiv, 0.1 increments) as measured by nMS. ....S32

**Supporting Figure S11.** Raw data nMS spectra showing PROTAC mediated formation of the full KRAS-ACBI3-CRL2$^{VHL}$-UBE2R1(C93K)-Ub complex. ....S33

**Supporting Table S12.** Observed *m/z* values and measured MWs for the full POI-PROTAC-E3-E2-Ub degradation complex with KRAS as the POI. ....S34

Recombinant production of CRL2$^{VHL}$ protein. ....S36

Recombinant production of KRAS protein....S37

**Supporting Table S13.** UHMR MS parameters employed for the methods used to characterise proteins and/or complexes. ....S39

References ....S41

**Table S1.** Sequences and calculated molecular weight (MW) of protein constructs utilised in this study.

| Protein | Subunit/s | Sequence | Subunit/s MW | Protein/protein complex MW |
|---|---|---|---|---|
| BRD4$^{BD2}$ (residues 333-460) | N/A | SMKDVPDSQQHPAPEKSSKVSEQLKCCSGILKEMF AKKHAAYAWPFYKPVDVEALGLHDYCDIIKHPMDM STIKSKLEAREYRDAQEFGADVRLMFSNCYKYNPP DHEVVAMARKLQDVFEMRFAKMPDE | N/A | 15,036.3 |
| KRAS (residues 1-169, GDP-bound form) | N/A | HMTEYKLVVVGAGGVGKSALTIQLIQNHFVDEYDP TIEDSYRKQVVIDGETCLLDILDTAGQEEYSAMRDQ YMRTGEGFLCVFAINNTKSFEDIHHYREQIKRVKDSE DVPMVLVGNKCDLPSRTVDTKQAQDLARSYGIPFIE TSAKTRQGVDDAFYTLVREIRKHKEK | N/A | 19,850.4[a] |
| VCB (VHL-EloC-EloB) | VHL (residues 54-213) | GSMEAGRPRPVLRSVNSREPSQVIFCNRSPRVVL PVWLNFDGEPQPYPTLPPGTGRRIHSYRGHLWLF RDAGTHDGLLVNQTELFVPSLNVDGQPIFANITLP VYTLKERCLQVVRSLVKPENYRRLDIVRSLYEDLE DHPNVQKDLERLTQERIAHQRMGD | 18,676.3 | 41,373.3 |
| | EloC (residues 1-104) | MDVFLMIRRHKTTIFTDAKESSTVFELKRIVEGILK RPPDEQRLYKDDQLLDDGKTLGECGFTSQTARP QAPATVGLAFRADDTFEALCIEPFSSPPELPDVMK | 11,733.4 | |
| | EloB (residues 17-112) | MMYVKLISSDGHEFIVKREHALTSGTIKAMLSGP GQFAENETNEVNFREIPSHVLSKVCMYFTYKVR YTNSSTEIPEFPIAPEIALELLMAANFLDC | 10,963.6 | |
| CRL2$^{VHL}$ (VHL-EloC-EloB-Cul2-Rbx1) | VHL | GSMEAGRPRPVLRSVNSREPSQVIFCNRSPRVVL PVWLNFDGEPQPYPTLPPGTGRRIHSYRGHLWLF | 18,676.3 | 140,831.5 |

| | | |
|---|---|---|
| (residues 54-213) | RDAGTHDGLLVNQTELFVPSLNVDGQPIFANITLP VYTLKERCLQVVRSLVKPENYRRLDIVRSLYEDLE DHPNVQKDLERLTQERIAHQRMGD | |
| EloC (residues 1-104) | MDVFLMIRRHKTTIFTDAKESSTVFELKRIVEGILK RPPDEQRLYKDDQLLDDGKTLGECGFTSQTARP QAPATVGLAFRADDTFEALCIEPFSSPPELPDVMK | 11,733.4 |
| EloB (residues 17-112) | MMYVKLISSDGHEFIVKREHALTSGTIKAMLSGP GQFAENETNEVNFREIPSHVLSKVCMYFTYKVR YTNSSTEIPEFPIAPEIALELLMAANFLDC | 10,963.6 |
| Cul2 (residues 1-745) | GGSMSLKPRVVDFDETWNKLLTTIKAVVMLEYVER ATWNDRFSDIYALCVAYPEPLGERLYTETKIFLENH VRHLHKRVLESEEQVLVMYHRYWEEYSKGADYMD CLYRYLNTQFIKKNKLTEADLQYGYGGVDMNEPLM EIGELALDMWRKLMVEPLQAILIRMLLREIKNDRGG EDPNQKVIHGVINSFVHVEQYKKKFPLKFYQEIFES PFLTETGEYYKQEASNLLQESNCSQYMEKVLGRLK DEEIRCRKYLHPSSYTKVIHECQQRMVADHLQFLH AECHNIIRQEKKNDMANMYVLLRAVSTGLPHMIQEL QNHIHDEGLRATSNLTQENMPTLFVESVLEVHGKF VQLINTVLNGDQHFMSALDKALTSVVNYREPKSVC KAPELLAKYCDNLLKKSAKGMTENEVEDRLTSFITV FKYIDDKDVFQKFYARMLAKRLIHGLSMSMDSEEA MINKLKQACGYEFTSKLHRMYTDMSVSADLNNKF NNFIKNQDTVIDLGISFQIYVLQAGAWPLTQAPSST FAIPQELEKSVQMFELFYSQHFSGRKLTWLHYLCT GEVKMNYLGKPYVAMVTTYQMAVLLAFNNSETVS YKELQDSTQMNEKELTKTIKSLLDVKMINHDSEKE DIDAESSFSLNMNFSSKRTKFKITTSMQKDTPQEM | 87,184.2 |

| | | | | |
|---|---|---|---|---|
| | | EQTRSAVDEDRKMYLQAAIVRIMKARKVLRHNALI QEVISQSRARFNPSISMIKKCIEVLIDKQYIERSQAS ADEYSYVA | | |
| | Rbx1 (residues 1-108) | MAAAMDVDTPSGTNSGAGKKRFEVKKWNAVALW AWDIVVDNCAICRNHIMDLCIECQANQASATSEEC TVAWGVCNHAFHFHCISRWLKTRQVCPLDNREW EFQKYGH | 12,274.0 | |
| UBE2R1 (C93K) | N/A | ARPLVPSSQKALLLELKGLQEEPVEGFRVTLVDE GDLYNWEVAIFGPPNTYYEGGYFKARLKFPIDYP YSPPAFRFLTKMWHPNIYETGDVKISILHPPVDDP QSGELPSERWNPTQNVRTILLSVISLLNEPNTFS PANVDASVMYRKWKESKGKDREYTDIIRKQVLG TKVDAERDGVKVPTTLAEYCVKTKAPAPDEGSD LFYDDYYEDGEVEEEADSCFGDDEDDSGTEES | N/A | 26,630.6 |
| UBE2RI (C93K)-$^{15}$N-Ub | UBE2R1 (C93K) | ARPLVPSSQKALLLELKGLQEEPVEGFRVTLVDE GDLYNWEVAIFGPPNTYYEGGYFKARLKFPIDYP YSPPAFRFLTKMWHPNIYETGDVKISILHPPVDDP QSGELPSERWNPTQNVRTILLSVISLLNEPNTFS PANVDASVMYRKWKESKGKDREYTDIIRKQVLG TKVDAERDGVKVPTTLAEYCVKTKAPAPDEGSD LFYDDYYEDGEVEEEADSCFGDDEDDSGTEES | 26,630.6 | 35,283.4[b] |
| | $^{15}$N-Ub | <u>M</u>QIFVKTLTGKTITLEVEPSDTIENVKAKIQDKEG IPPDQQRLIFAGKQLEDGRTLSDYNIQKESTLHL VLRLRGG | 8,670.8 | |

[a]Expected MW calculated considers the protein sequence (19,382.9 Da) and bound Guanosine 5′-diphosphate (GDP, 443.2 Da) and $Mg^{2+}$ (24.3 Da). [b]Expected MW calculated considers the covalent bond between E2 and Ub (Δ -18 Da).

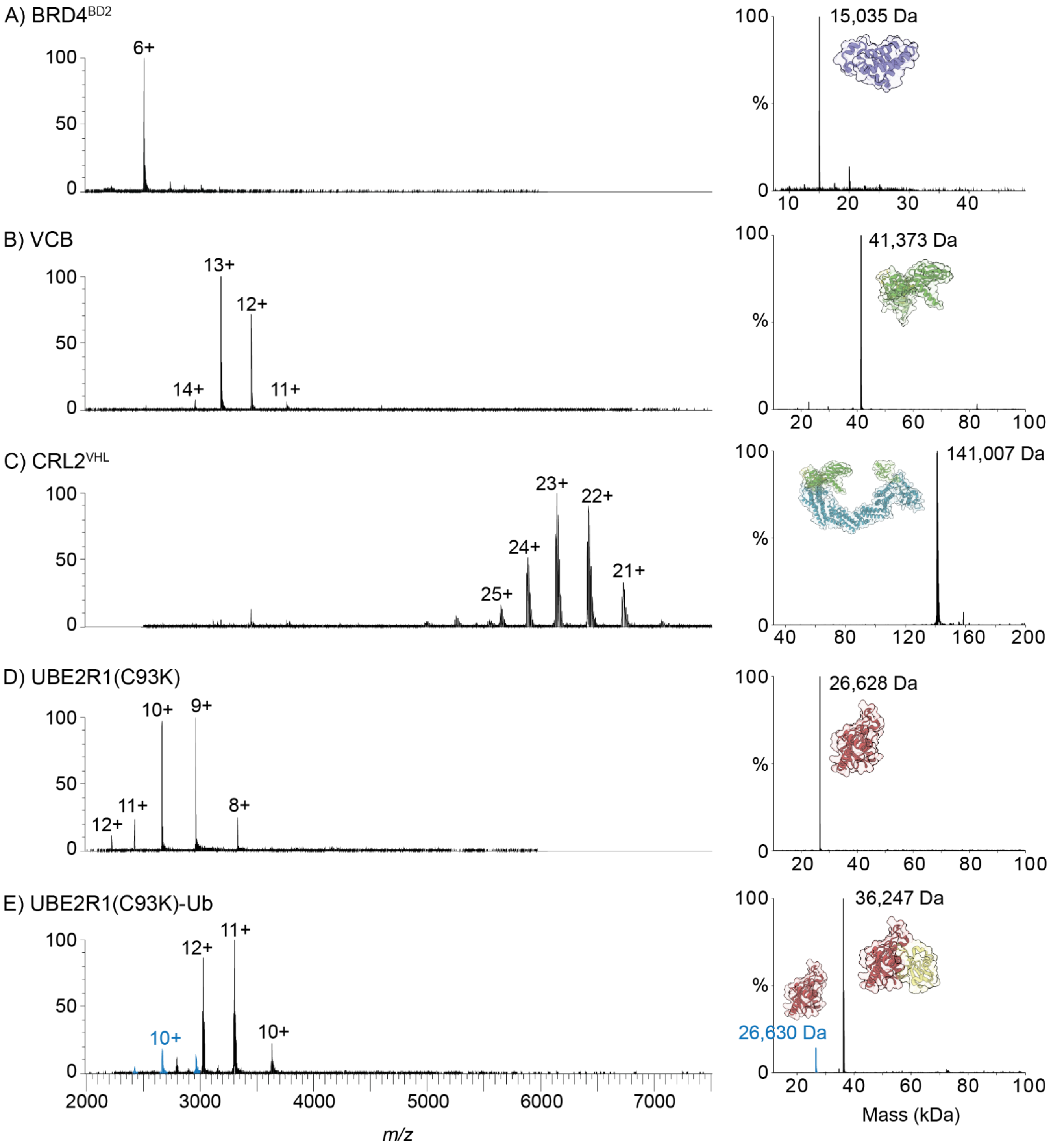

**Supporting Figure S1.** Raw and deconvoluted nMS spectra for individual protein components (150 mM $NH_4OAc$). A) 1 µM BRD4$^{BD2}$, expected MW is 15,036 Da. B) 1 µM VCB, expected MW is 41,373 Da. C) 1.5 µM CRL2$^{VHL}$, expected MW is 140,832 Da (or ~141,023 Da with 3× $Zn^{2+}$ cations as observed in PDB 8RWX and PDB 8RX0). D) 1 µM UBE2R1(C93K), expected MW is 26,631 Da. E) 1 µM UBE2R1(C93K)-Ub, expected MW is 35,283Da ($^{15}$N-labeled Ub is used, referred to as 'Ub' throughout remaining analysis). The blue peaks and labels in E) correspond to a minor (~9.6%) amount of E2 with no Ub conjugated. Observed *m/z* values and MWs in **Supporting Tables S2** and **S3**.

**Supporting Table S2.** Observed *m/z* values, charge states and measured MWs for individual protein components. Raw and deconvoluted mass spectra are shown in **Supporting Figure S1**.

| **Protein Component** | **Expected MW (Da)** | **Observed *m/z*** | **Charge state (*z*)** | **Deconvoluted MW (Da)[a]** |
|---|---|---|---|---|
| BRD4$^{BD2}$ | 15,036.3 | 2506.8765 | 6+ | 15,035 |
| VCB (VHL-EloC-EloB) | 41,373.4 | 2956.3498 | 14+ | 41,373 |
| | | 3183.5237 | 13+ | |
| | | 3448.6302 | 12+ | |
| | | 3762.0383 | 11+ | |
| CRL2$^{VHL}$ (VHL-EloC-EloB-Cul2-Rbx1) | 140,831.5 (or ~141,023 Da with 3× $Zn^{2+}$ cations as observed in PDB 8RWX and PDB 8RX0) | 5641.4632 | 25+ | 141,007 |
| | | 5876.6072 | 24+ | |
| | | 6132.4421 | 23+ | |
| | | 6411.3233 | 22+ | |
| | | 6716.9335 | 21+ | |
| | | 7052.9235 | 20+ | |
| UBE2R1(C93K) | 26,630.6 | 2219.9264 | 12+ | 26,628 |
| | | 2421.7416 | 11+ | |
| | | 2663.818 | 10+ | |
| | | 2959.6957 | 9+ | |
| | | 3329.5262 | 8+ | |
| UBE2R1(C93K)-$^{15}$N-Ub | 35,283.5[b] | 2421.8754 | 11+ | 26,630[c] |
| | | 2663.9924 | 10+ | |
| | | 2959.8884 | 9+ | |
| | | 2789.2336 | 13+ | 36,247 |
| | | 3021.6625 | 12+ | |
| | | 3296.2424 | 11+ | |
| | | 3625.7357 | 10+ | |

[a]MW deconvoluted in UniDec,[2] to nearest whole Da. [b]Expected MW calculated considers the covalent bond between E2 and Ub (Δ -18 Da). [c]Corresponds to a minor (~9.6%) amount of E2 with no Ub conjugated.

**Supporting Table S3.** Reference corrected MW of protein components, PROTAC/compound MWs and expected MW of protein complexes determined (based on corrected MW).

| **Protein** | **Corrected MW (Da)**[a] |
|---|---|
| BRD4$^{BD2}$ | 15,036[b] |
| VCB | 41,373[b] |
| CRL2$^{VHL}$ | 141,007[c] |
| UBE2R1(C93K) | 26,631[b] |
| UBE2R1(C93K)-Ub | 36,247[c] |
| **Compounds/PROTACs** | **MW (Da)** |
| MZ1 | 1002.6 |
| *cis*-MZ1 | 1002.6 |
| **Protein Complexes** | **Expected MW (Da)**[d] |
| BRD4$^{BD2}$-VCB[e] | 56,409 |
| BRD4$^{BD2}$-MZ1-VCB[f] | 57,412 |
| BRD4$^{BD2}$-CRL2$^{VHL}$[e] | 156,043 |
| BRD4$^{BD2}$-MZ1-CRL2$^{VHL}$[f] | 157,046 |
| CRL2$^{VHL}$-UBE2R1(C93K) | 167,638 |
| CRL2$^{VHL}$-UBE2R1(C93K)-Ub | 177,254 |
| BRD4$^{BD2}$-CRL2$^{VHL}$-UBE2R1(C93K)-Ub[e] | 192,290 |
| BRD4$^{BD2}$-MZ1-CRL2$^{VHL}$-UBE2R1(C93K)-Ub[f] | 193,293 |

[a]Corrected MW is based on the nMS analysis for protein components (**Supporting Figure S1**) and are used for data analysis throughout the rest of the study. [b]MW as determined from protein sequence (corresponds with MW as measured by nMS, **Supporting Figure S1**). [c]MW as determined from nMS visibility studies (**Supporting Figure S1**) due to differences with that expected from protein sequence. [d]Expected MWs of protein complexes of interest for this study, calculated using the corrected MWs for individual protein components (rounded to nearest Da). [e]Complexes without PROTAC and not necessarily expected to form, but MWs provided to compare with spectra to confirm presence or absence. [f]Complexes containing *cis*-MZ1 are not expected to form, however, due to the MW of *cis*-MZ1, any such complexes would have the same MW as the MZ1-containing complexes.

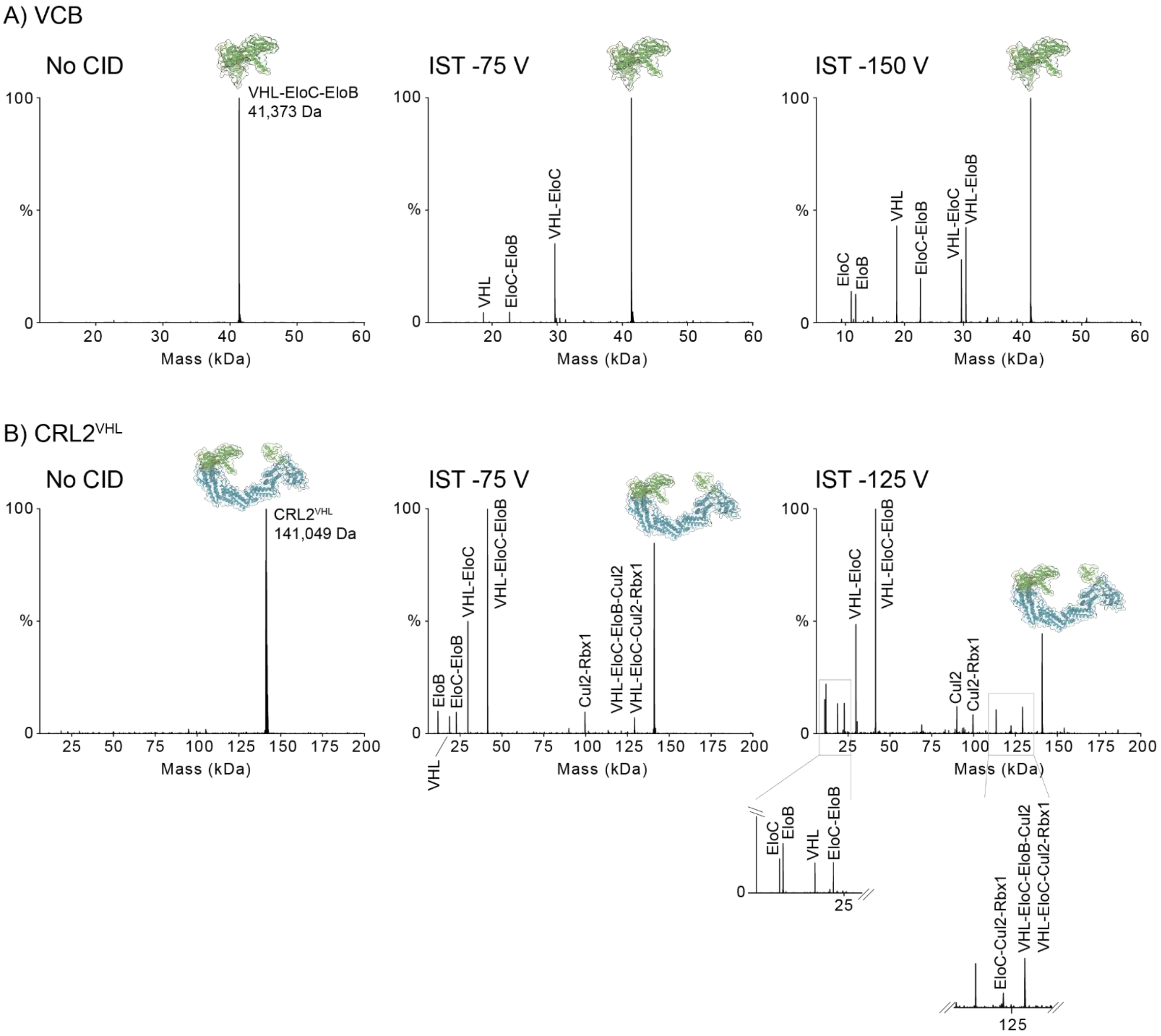

**Supporting Figure S2.** Collision-induced dissociation (CID) of minimal E3 (VCB) and full E3 (CRL2$^{VHL}$) into individual component subunits. A) VCB (1 µM) analysis without CID and with subunit dissociation via in-source trapping desolvation voltages -75 V and -150 V. B) CRL2$^{VHL}$ (1 µM) analysis without CID and with subunit dissociation via in-source trapping desolvation voltages -75 V and -125 V (CRL2$^{VHL}$ was not visible at -150 V for a direct comparison with VCB analysis). Deconvoluted spectra are annotated with the dissociated component identities; unlabelled peaks were unable to be assigned to a known intact subunit/s. Observed MWs are provided in **Supporting Table S4.**

**Supporting Table S4.** Observed MW and assignment to subunits dissociated from E3s VCB and CRL2$^{VHL}$ with in-source CID. Deconvoluted mass spectra are in **Supporting Figure S2**.

| **Protein** | **CID V** | **Figure/Spectra** | **Observed MW (Da)**$^{a}$ | **Assigned Subunits** | **Expected MW (Da)** |
|---|---|---|---|---|---|
| VCB | 0 | S2A, left | 41,373 | VHL-EloC-EloB | 41,373 |
| | -75 | S2A, middle | 18,675 | VHL | 18,676 |
| | | | 22,696 | EloC-EloB | 22,697 |
| | | | 29,639 | VHL-EloC | 29,640 |
| | | | 41,373 | VHL-EloC-EloB | 41,373 |
| | -150 | S2A, right | 10,962 | EloC | 10,964 |
| | | | 11,732 | EloB | 11,733 |
| | | | 18,675 | VHL | 18,676 |
| | | | 22,695 | EloC-EloB | 22,697 |
| | | | 29,639 | VHL-EloC | 29,640 |
| | | | 30,408 | VHL-EloB | 30,410 |
| | | | 41,372 | VHL-EloC-EloB | 41,373 |
| CRL2$^{VHL}$ | 0 | S2B, left | 141,049 | VHL-EloC-EloB-Cul2-Rbx1 | 140,832 |
| | -75 | S2B, middle | 11,732 | EloB | 11,733 |
| | | | 18,676 | VHL | 18,676 |
| | | | 22,697 | EloC-EloB | 22,697 |
| | | | 29,638 | VHL-EloC | 29,640 |
| | | | 41,369 | VHL-EloC-EloB | 41,373 |
| | | | 99,583 | Cul2-Rbx1 | 99,458 |
| | | | 129,257 | VHL-EloC-EloB-Cul2 or VHL-EloC-Cul2-Rbx1 | 128,558 or 129,098 |
| | | | 141,000 | VHL-EloC-EloB-Cul2-Rbx1 | 140,832 |
| | -125 | S2B, right | 10,962 | EloC | 10,964 |
| | | | 11,731 | EloB | 11,733 |
| | | | 18,676 | VHL | 18,676 |
| | | | 22,696 | EloC-EloB | 22,697 |
| | | | 29,637 | VHL-EloC | 29,640 |
| | | | 30,408 | VHL-EloB | 30,410 |
| | | | 41,370 | VHL-EloC-EloB | 41,373 |
| | | | 68,996 | [b, c] | |
| | | | 89,954 | Cul2 | 87,184 |
| | | | 99,575 | Cul2-Rbx1 | 99,458 |
| | | | 113,455 | [b] | |
| | | | 122,365 | EloC-EloB-Cul2-Rbx1[c] | 122,155 |
| | | | 129,239 | VHL-EloC-EloB-Cul2 or VHL-EloC-Cul2-Rbx1 | 128,558 or 129,098 |
| | | | 140,971 | VHL-EloC-EloB-Cul2-Rbx1 | 140,832 |

[a]MW deconvoluted in UniDec,[2] to nearest whole Da. [b]Observed MW unable to be assigned to a known intact subunit/s. [c]Protein complexes were observed at relative intensity <5%.

**Supporting Table S5.** Observed *m/z* values and measured MWs for binary binding of MZ1 (MW 1002.6 Da) and *cis*-MZ1 (MW 1002.6 Da) to BRD4$^{BD2}$, VCB and CRL2$^{VHL}$. Raw mass spectra (of the most intense charge states) are shown in **Figure 3**. Binary complexes – blue text.

| **Sample** | | **Protein Species** | **Expected MW (Da)** | **Observed *m/z*** | **Charge State (*z*)** | **Deconvoluted MW (Da)**$^{a}$ | **Bound Ligand MW (ΔMW) (Da)** |
|---|---|---|---|---|---|---|---|
| **Protein** | **Ligand** | | | | | | |
| BRD4$^{BD2}$ | - | BRD4$^{BD2}$ | 15,036.3 | 2506.8596<br>3007.8315 | 6+<br>5+ | 15,035 | N/A |
| | MZ1 | BRD4$^{BD2}$ | 15,036.3 | 2506.6916<br>3008.0313 | 6+<br>5+ | 15,034 | N/A |
| | | BRD4$^{BD2}$-MZ1 | 16,038.9 | 2292.2146<br>2674.2514 | 7+<br>6+ | 16,038 | 1,002 |
| | *cis*-MZ1 | BRD4$^{BD2}$ | 15,036.3 | 2507.0247<br>3008.4322 | 6+<br>5+ | 15,036 | N/A |
| | | BRD4$^{BD2}$-*cis*-MZ1 | 16,038.9 | 2291.9269<br>2674.0825 | 7+<br>6+ | 16,038 | 1,002 |
| VCB | - | VCB | 41,373.4 | 3448.5599<br>3762.1009<br>4138.1978<br>4597.819 | 12+<br>11+<br>10+<br>9+ | 41,373 | N/A |
| | MZ1 | VCB | 41,373.4 | 3448.5423<br>3762.0871<br>4138.1848<br>4597.8366 | 12+<br>11+<br>10+<br>9+ | 41,373 | N/A |
| | | VCB-MZ1 | 42,376.0 | 3532.2241<br>3853.1851<br>4238.3755<br>4709.4314 | 12+<br>11+<br>10+<br>9+ | 42,375 | 1,002 |
| | *cis*-MZ1 | VCB | 41,373.4 | 3448.622<br>3762.1614<br>4138.2476<br>4597.9403 | 12+<br>11+<br>10+<br>9+ | 41,373 | N/A |
| CRL2$^{VHL}$ | - | CRL2$^{VHL}$ | 141,007 | 6412.0768<br>6717.8703<br>7053.887<br>7425.1089<br>7837.7589 | 22+<br>21+<br>20+<br>19+<br>18+ | 141,060 | N/A |
| | MZ1 | CRL2$^{VHL}$ | 141,007 | 6411.0288<br>6719.9109<br>7053.098<br>7424.2263 | 22+<br>21+<br>20+<br>19+ | 141,039 | N/A |

| | | | | | | |
|---|---|---|---|---|---|---|
| | | | 7837.1655 | 18+ | | |
| | CRL2$^{VHL}$-MZ1 | 142,010 | 6456.9744 | 22+ | 142,035 | 996 |
| | | | 6764.1866 | 21+ | | |
| | | | 7103.0769 | 20+ | | |
| | | | 7476.8274 | 19+ | | |
| | | | 7892.6892 | 18+ | | |
| *cis*-MZ1 | CRL2$^{VHL}$ | 141,007 | 6411.3432 | 22+ | 141,042 | N/A |
| | | | 6717.0904 | 21+ | | |
| | | | 7053.5664 | 20+ | | |
| | | | 7424.9112 | 19+ | | |
| | | | 7837.5369 | 18+ | | |

[a]MW deconvoluted in UniDec,[2] to nearest whole Da and with relative intensity >5-10%. N/A: not applicable as no ligand binding observed.

**Supporting Table S6.** Observed *m/z* values and measured MWs for samples comprising BRD4[BD2] and either VCB or CRL2[VHL] with MZ1 (MW 1002.6 Da) and *cis*-MZ1 (MW 1002.6 Da). Raw mass spectra as shown in **Figure 4**. Binary complexes – blue text, ternary complexes – red text.

| Sample POI | E3 | Compound | Observed Species | Expected MW (Da) | Observed *m/z* | Charge State (*z*) | Deconvoluted MW (Da)[a] |
|---|---|---|---|---|---|---|---|
| BRD4[BD2] | VCB | - | BRD4[BD2] | 15,036.3 | 2506.7104 | 6+ | 15,034 |
| | | | VCB | 41,373.4 | 3448.6663 | 12+ | 41,372 |
| | | | | | 3762.116 | 11+ | |
| | | | | | 4138.1691 | 10+ | |
| | | | | | 4597.8007 | 9+ | |
| | | MZ1 | BRD4[BD2] | 15,036.3 | 2506.6251 | 6+ | [b] |
| | | | BRD4[BD2]-MZ1 | 16,038.9 | 2292.1765 | 7+ | [b] |
| | | | | | 2673.8921 | 6+ | |
| | | | VCB | 41,373.4 | 3448.5386 | 12+ | 41,372 |
| | | | | | 3762.0472 | 11+ | |
| | | | | | 4138.1602 | 10+ | |
| | | | VCB-MZ1 | 42,376.0 | 3532.1979 | 12+ | 42,374 |
| | | | | | 3853.1694 | 11+ | |
| | | | | | 4238.3828 | 10+ | |
| | | | BRD4[BD2]-MZ1-VCB | 57,412.3 | 3828.3368 | 15+ | 57,409 |
| | | | | | 4101.6456 | 14+ | |
| | | | | | 4417.0471 | 13+ | |
| | | | | | 4784.988 | 12+ | |
| | | | | | 5220.0705 | 11+ | |
| | | *cis*-MZ1 | BRD4[BD2] | 15,036.3 | 2506.6498 | 6+ | 15,037 |
| | | | BRD4[BD2]-*cis*-MZ1 | 16,038.9 | 2291.9968 | 7+ | 16,038 |
| | | | | | 2673.8488 | 6+ | |
| | | | VCB | 41,373.4 | 3448.6715 | 12+ | 41,372 |
| | | | | | 3762.1097 | 11+ | |
| | | | | | 4138.1529 | 10+ | |
| | | | | | 4597.7988 | 9+ | |
| | CRL2[VHL] | - | BRD4[BD2] | 15,036.3 | 2149.0475 | 7+ | 15,036 |
| | | | | | 2506.8536 | 6+ | |
| | | | | | 3008.0689 | 5+ | |
| | | | VCB | 41,373.4 | 3448.6337 | 12+ | 41,371 |
| | | | | | 3761.9333 | 11+ | |
| | | | | | 4138.4009 | 10+ | |
| | | | | | 4598.1230 | 9+ | |
| | | | Cul2-Rbx1 | 99,458.2 | 5533.3442 | 18+ | 99,590 |
| | | | | | 5859.2324 | 17+ | |
| | | | | | 6225.5383 | 16+ | |
| | | | | | 6640.5519 | 15+ | |
| | | | CRL2[VHL] | 141,007 | 6409.2994 | 22+ | 140,996 |

| | | | | | |
|---|---|---|---|---|---|
| | | | 6714.8612 | 21+ | |
| | | | 7050.951 | 20+ | |
| | | | 7422.4237 | 19+ | |
| | | | 7835.0047 | 18+ | |
| | | | 8297.3235 | 17+ | |
| MZ1 | BRD4$^{BD2}$ | 15,036.3 | 2148.877 | 7+ | 15,036 |
| | | | 2506.8293 | 6+ | |
| | VCB | 41,373.4 | 3448.6723 | 12+ | 41,369 |
| | | | 3761.8423 | 11+ | |
| | | | 4137.836 | 10+ | |
| | | | 4597.8113 | 9+ | |
| | BRD4$^{BD2}$-VCB | 56,409.7 | 4340.2257 | 13+ | 56,411 |
| | | | 4701.9928 | 12+ | |
| | | | 5129.4489 | 11+ | |
| | BRD4$^{BD2}$-MZ1-VCB | 57,412.3 | 4417.4739 | 13+ | 57,415 |
| | | | 4785.5243 | 12+ | |
| | | | 5220.6368 | 11+ | |
| | Cul2-Rbx1 | 99,458.2 | 5533.2309 | 18+ | 99,580 |
| | | | 5858.6474 | 17+ | |
| | | | 6224.9399 | 16+ | |
| | | | 6639.9786 | 15+ | |
| | CRL2$^{VHL}$ | 141,007.0 | 6409.344 | 22+ | 140,999 |
| | | | 6714.964 | 21+ | |
| | | | 7051.527 | 20+ | |
| | | | 7422.7996 | 19+ | |
| | | | 7835.0374 | 18+ | |
| | BRD4$^{BD2}$-MZ1-CRL2$^{VHL}$ | 157,045.6 | 6543.2716 | 24+ | 157,069 |
| | | | 6828.7287 | 23+ | |
| | | | 7140.0304 | 22+ | |
| | | | 7480.2482 | 21+ | |
| | | | 7854.3152 | 20+ | |
| | | | 8268.4585 | 19+ | |
| | | | 8728.082 | 18+ | |
| *cis*-MZ1 | BRD4$^{BD2}$ | 15,036.3 | 2157.5124 | 7+ | 15,035 |
| | | | 2506.8337 | 6+ | |
| | | | 3008.1005 | 5+ | |
| | VCB | 41,373.4 | 3448.6355 | 12+ | 41,373 |
| | | | 3762.1336 | 11+ | |
| | | | 4138.336 | 10+ | |
| | Cul2-Rbx1 | 99,458.2 | 5534.6643 | 18+ | 99,611 |
| | | | 5860.1552 | 17+ | |
| | | | 6227.1531 | 16+ | |
| | CRL2$^{VHL}$ | 141,007 | 6411.263 | 22+ | 141,038 |
| | | | 6716.9436 | 21+ | |
| | | | 7052.8503 | 20+ | |
| | | | 7424.2839 | 19+ | |

| | | |
|---|---|---|
| | 7837.3589 | 18+ |
| | 8299.3488 | 17+ |

[a]MW deconvoluted in UniDec,[2] to nearest whole Da and with relative intensity >5%. [b]Species not detected during deconvolution but evident in raw spectrum.

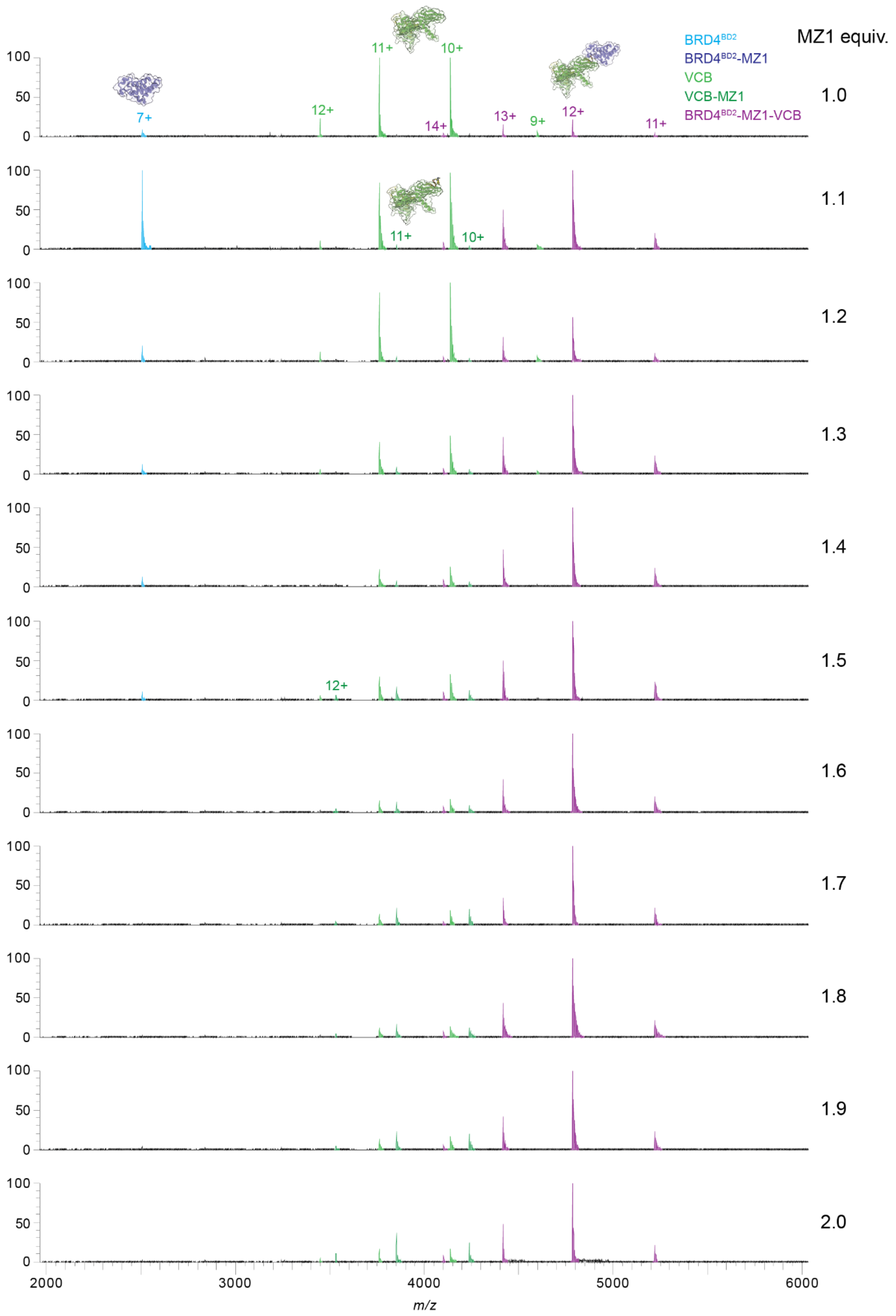

**Supporting Figure S3.** nMS titration of PROTAC MZ1 (1.0-2.0 equiv, 0.1 increments) against POI BRD4$^{BD2}$ and E3 VCB (present at equimolar concentrations, 1 μM each) to determine the optimal MZ1 equiv for maximal ternary complex formation.

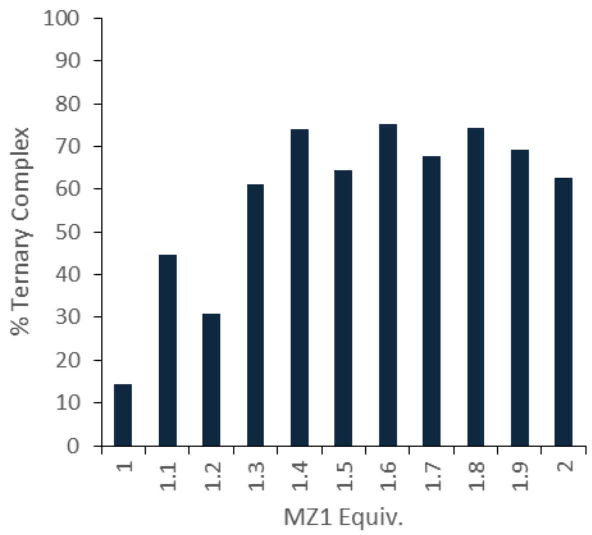

**Supporting Figure S4.** Quantification of ternary complex formation between $BRD4^{BD2}$ and VCB (present at equimolar concentrations, 1 µM each) with MZ1 titrated (1.0-2.0 equiv, 0.1 increments) as measured by nMS. The % ternary complex formed was calculated from the deconvoluted data of mass spectra in **Supporting Figure S3** as the ratio of VCB recruited into the ternary complex (i.e. sum of the intensity of the deconvoluted ternary complex peak divided by the sum of the intensities of the deconvoluted peaks for unbound VCB, MZ1-bound VCB and ternary complex).

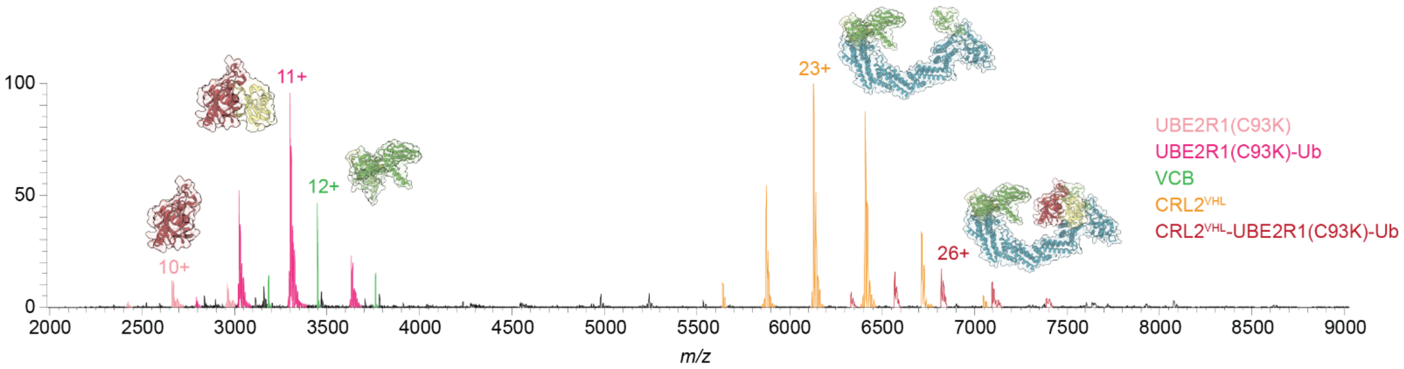

**Supporting Figure S5.** Raw data nMS spectra of E3-E2-Ub sample comprising CRL2$^{VHL}$ (1 µM) and UBE2R1(C93K)-Ub (5 µM). E3-E2-Ub complex (CRL2$^{VHL}$-UBE2R1(C93K)-Ub, red) was detected, along with free CRL2$^{VHL}$ (orange), dissociated VCB (green), free UBE2R1(C93K)-Ub (dark pink, and contaminating UBE2R1(C93K) without Ub conjugated, light pink). Where the deconvoluted species was present at >5% relative intensity the peaks are labelled, including with the charge state for the most intense peak for each observed species. Corresponding deconvoluted nMS are in manuscript **Figure 5A** and observed *m/z* values and MWs in **Supporting Table S7**

**Supporting Table S7.** Observed *m/z* values, charge states and measured MWs for species observed in the nMS spectra of the E3-E2-Ub sample (CRL2$^{VHL}$ and UBE2R1(C93K)-Ub). Raw and deconvoluted mass spectra are shown in **Supporting Figure S5** and **Figure 5A**, respectively.

| **Sample** | **Protein Species** | **Expected MW (Da)** | **Observed *m/z*** | **Charge State (z)** | **Deconvoluted MW (Da)**[a] |
|---|---|---|---|---|---|
| CRL2$^{VHL}$, UBE2R1(C93K)-Ub | UBE2R1(C93K) | 26,630.6 | 2421.7311 | 11+ | 26,630 |
| | | | 2664.0343 | 10+ | |
| | | | 2959.9475 | 9+ | |
| | UBE2R1(C93K)-Ub | 36,247.0 | 2789.265 | 13+ | 36,308 |
| | | | 3021.7972 | 12+ | |
| | | | 3296.3306 | 11+ | |
| | | | 3625.7973 | 10+ | |
| | VHL-EloC-EloB | 41,373.4 | 3183.5315 | 13+ | 41,370 |
| | | | 3448.4197 | 12+ | |
| | | | 3762.0578 | 11+ | |
| | CRL2$^{VHL}$ | 141,007.0 | 5639.5119 | 25+ | 140,969 |
| | | | 5874.7173 | 24+ | |
| | | | 6130.2455 | 23+ | |
| | | | 6409.0395 | 22+ | |
| | | | 6713.9964 | 21+ | |
| | | | 7049.544 | 20+ | |
| | CRL2$^{VHL}$-UBE2R1(C93K)-Ub | 177,254.0 | 6333.9635 | 28+ | 177,339 |
| | | | 6568.7772 | 27+ | |
| | | | 6821.5043 | 26+ | |
| | | | 7095.0852 | 25+ | |
| | | | 7390.1709 | 24+ | |

[a]MW deconvoluted in UniDec,[2] to nearest whole Da and with relative intensity >5%.

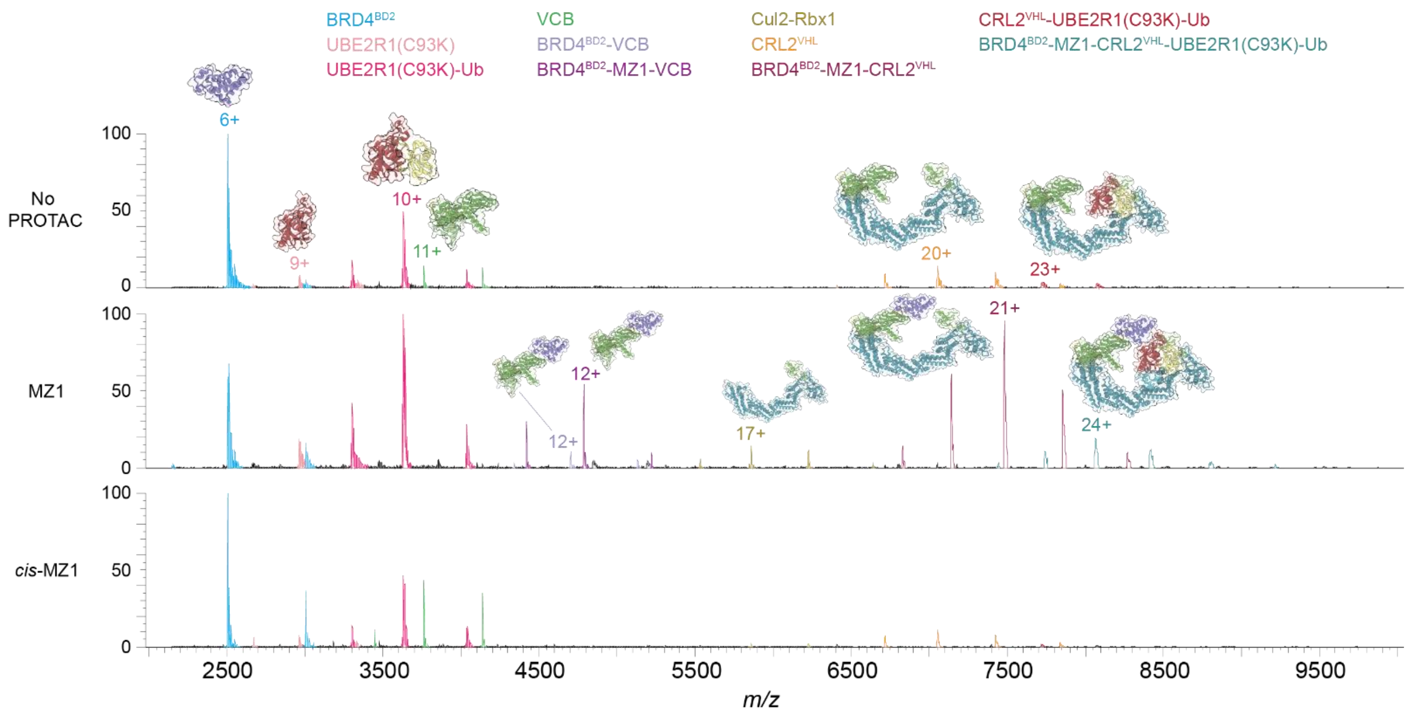

**Supporting Figure S6.** Raw data nMS spectra showing PROTAC mediated formation of the full BRD4$^{BD2}$-MZ1-CRL2$^{VHL}$-UBE2R1(C93K)-Ub complex. The peaks for each observed species that were present at >3% relative intensity following deconvolution are labelled via the indicated colour coding, and the charge state for the most intense peak for each observed species is also labelled. Corresponding deconvoluted nMS are shown in **Figure 5B-D** and observed *m/z* values and MWs provided in **Supporting Table S8**.

**Supporting Table S8.** Observed *m/z* values and measured MWs for the full POI-PROTAC-E3-E2-Ub degradation complex with BRD4$^{BD2}$ as the POI. Raw and deconvoluted mass spectra are shown in **Supporting Figure S6** and **Figure 5B-D**, respectively). Binary complexes – blue text, ternary complexes – red text, full complex in green text.

| Sample Proteins | Compound | Protein Species | Expected MW (Da) | Observed *m/z* | Charge State (z) | Deconvoluted MW (Da)$^{a}$ |
|---|---|---|---|---|---|---|
| BRD4$^{BD2}$, CRL2$^{VHL}$, UBE2R1(C93K)-Ub | - | BRD4$^{BD2}$ | 15,036.3 | 2506.9248<br>3008.4526 | 6+<br>5+ | 15,036 |
| | | UBE2R1(C93K) | 26,630.6 | 2669.9870<br>2959.9446<br>3337.5135 | 10+<br>9+<br>8+ | 26,631 |
| | | UBE2R1(C93K)-Ub | 36,247.0 | 3301.6857<br>3631.8879<br>4035.2731 | 11+<br>10+<br>9+ | 36,308 |
| | | VHL-EloB-EloC | 41,373.4 | 3762.2789<br>4138.4433 | 11+<br>10+ | 41,374 |
| | | CRL2$^{VHL}$ | 141,007.0 | 6409.1024<br>6715.4128<br>7051.9532<br>7424.0482<br>7838.6688 | 22+<br>21+<br>20+<br>19+<br>18+ | 141,020 |
| | | CRL2$^{VHL}$-UBE2R1(C93K)-Ub | 177,254.0 | 7721.0020<br>8072.8616 | 23+<br>22+ | 177,589 |
| | MZ1 | BRD4$^{BD2}$ | 15,036.3 | 2149.0679<br>2506.8830<br>3008.4415 | 7+<br>6+<br>5+ | 15,037 |
| | | UBE2R1(C93K) | 26,630.6 | 2959.9261 | 9+ | 26,629 |
| | | UBE2R1(C93K)-Ub | 36,247.0 | 3301.6631<br>3631.7703<br>4035.2154 | 11+<br>10+<br>9+ | 36,308 |
| | | BRD4$^{BD2}$-VHL-EloC-EloB | 56,409.7 | 4340.2769<br>4702.2645<br>5129.5862 | 13+<br>12+<br>11+ | 56,415 |
| | | BRD4$^{BD2}$-MZ1-VHL-EloC-EloB | 57,412.3 | 4417.6583<br>4785.8801<br>5220.9530 | 13+<br>12+<br>11+ | 57,417 |
| | | Cul2-Rbx1 | 99,458.2 | 5533.2471<br>5858.9179<br>6225.1277<br>6716.1190 | 18+<br>17+<br>16+<br>15+ | 99,583 |

| | | | | | |
|---|---|---|---|---|---|
| | BRD4$^{BD2}$-MZ1-CRL2$^{VHL}$ | 157,045.9 | 6829.0166 | 23+ | 157,070 |
| | | | 7140.0173 | 22+ | |
| | | | 7480.3888 | 21+ | |
| | | | 7855.3868 | 20+ | |
| | | | 8269.2690 | 19+ | |
| | BRD4$^{BD2}$-MZ1-CRL2$^{VHL}$-UBE2R1(C93K)-Ub | 193,292.9 | 7446.4773 | 26+ | 193,479 |
| | | | 7738.4118 | 25+ | |
| | | | 8064.4410 | 24+ | |
| | | | 8419.5551 | 23+ | |
| | | | 8795.9065 | 22+ | |
| | | | 9217.7988 | 21+ | |
| *cis*-MZ1 | BRD4$^{BD2}$ | 15,036.3 | 2506.9647 | 6+ | 15,036 |
| | | | 3008.1412 | 5+ | |
| | UBE2R1(C93K) | 26,630.6 | 2959.9504 | 9+ | 26,630 |
| | UBE2R1(C93K)-Ub | 36,247.0 | 3301.7445 | 11+ | 36,310 |
| | | | 3631.9606 | 10+ | |
| | | | 4035.5041 | 9+ | |
| | VHL-EloB-EloC | 41,373.4 | 3448.6484 | 12+ | 41,372 |
| | | | 3761.9328 | 11+ | |
| | | | 4138.4838 | 10+ | |
| | Cul2-Rbx1 | 99,458.2 | 5532.6504 | 18+ | 99,584 |
| | | | 5858.6977 | 17+ | |
| | | | 6225.2509 | 16+ | |
| | CRL2$^{VHL}$ | 141,007.0 | 6409.3917 | 22+ | 141,038 |
| | | | 6715.605 | 21+ | |
| | | | 7052.3944 | 20+ | |
| | | | 7424.4538 | 19+ | |
| | | | 7837.0402 | 18+ | |
| | CRL2$^{VHL}$-UBE2R1(C93K)-Ub | 177,254.0 | 7399.0569 | 24+ | 177,465 |
| | | | 7715.8799 | 23+ | |
| | | | 8067.6858 | 22+ | |
| | | | 8452.0352 | 21+ | |

[a]MW deconvoluted in UniDec,[2] to nearest whole Da and with relative intensity >3%.

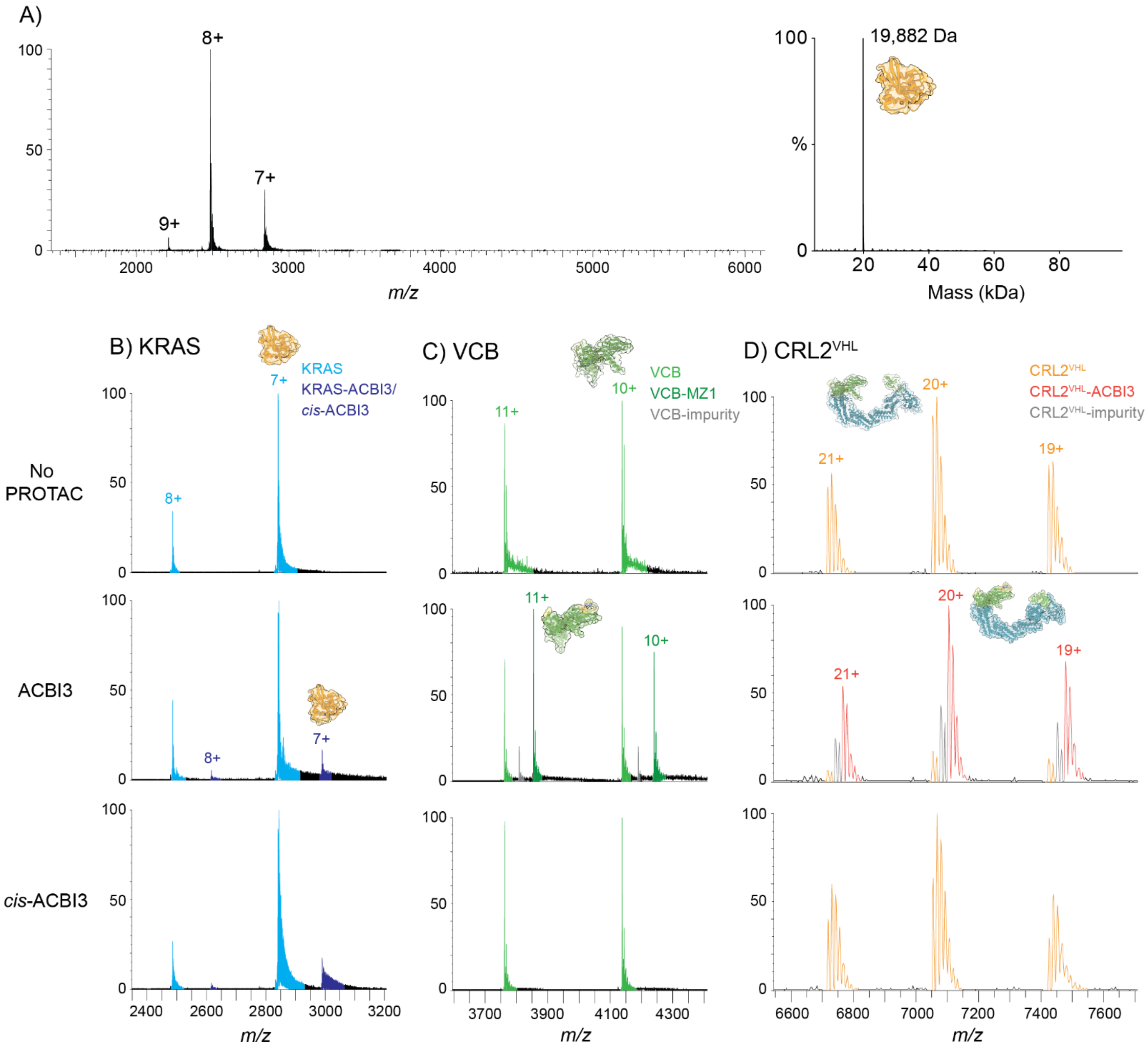

**Supporting Figure S7.** nMS assessment of KRAS and binary binding of ACBI3 and *cis*-ACBI3. A) Raw (LHS) and deconvoluted (RHS) nMS spectra (150 mM $NH_4OAc$) for 1 µM KRAS, expected MW is 19,850.4 Da, observed MW is 19,882 Da. B-D) Assessment of binary binding of PROTAC ACBI3 and negative control compound *cis*-ACBI3 (2 µM, 2 equiv) to 1 µM B) KRAS, C) VCB and D) CRL2$^{VHL}$ (150 mM $NH_4OAc$, 1% DMSO). An impurity in the ACBI3 sample (MW ~515 Da) was also observed binding to VCB and CRL2$^{VHL}$ but not to KRAS. The observed MW of 19,882 Da for KRAS is used for data analysis throughout the rest of the study. Observed *m/z* values and MWs are in **Supporting Table S9.**

**Supporting Table S9.** Observed *m/z* values and measured MWs for binary binding of ACBI3 (MW 1019.3 Da) and *cis*-ACBI3 (MW 1019.3 Da) to KRAS, VCB and CRL2$^{VHL}$. Raw spectra are shown in **Supporting Figure S7**. Binary complexes – blue text.

| Sample | | Protein Species | Expected MW (Da) | Observed *m/z* | Charge State (z) | Deconvoluted MW (Da)[a] | Bound Ligand MW (ΔMW) (Da) |
|---|---|---|---|---|---|---|---|
| **Protein** | **Ligand** | | | | | | |
| KRAS | - | KRAS | 19,882.0 | 2486.2000<br>2841.3773<br>3314.6055 | 8+<br>7+<br>6+ | 19,882 | N/A |
| | ACBI3 | KRAS | 19,882.0 | 2486.4516<br>2841.3788<br>3318.2690 | 8+<br>7+<br>6+ | 19,883 | N/A |
| | | KRAS-ACBI3 | 20,901.3 | 2616.2527<br>2990.0114 | 8+<br>7+ | 20,924[b] | 1,041[b] |
| | *cis*-ACBI3 | KRAS | 19,882.0 | 2486.1989<br>2841.236<br>3318.5996 | 8+<br>7+<br>6+ | 19,904[b] | N/A |
| | | KRAS-*cis*-ACBI3 | 20,901.3 | 2616.5008<br>2990.0069<br>3488.1714 | 8+<br>7+<br>6+ | 20,923[b] | 1,019 |
| VCB | - | VCB | 41,373.4 | 3448.7130<br>3762.0868<br>4138.0160<br>4596.8628 | 12+<br>11+<br>10+<br>9+ | 41,372 | N/A |
| | ACBI3 | VCB | 41,373.4 | 3448.5737<br>3762.0069<br>4138.1159<br>4597.9829 | 12+<br>11+<br>10+<br>9+ | 41,372 | N/A |
| | | VCB-impurity[c] | N/A | 3491.6228<br>3808.9210<br>4189.6719 | 12+<br>11+<br>10+ | 41,887 | 515[c] |
| | | VCB-ACBI3 | 42,392.7 | 3533.5743<br>3854.7044<br>4240.0195<br>4711.4084 | 12+<br>11+<br>10+<br>9+ | 42,392 | 1,020 |
| | *cis*-ABCI3 | VCB | 41,373.4 | 3448.5666<br>3762.0910<br>4138.1639<br>4597.7962 | 12+<br>11+<br>10+<br>9+ | 41,373 | N/A |
| CRL2$^{VHL}$ | - | CRL2$^{VHL}$ | 141,007 | 6412.0768<br>6717.8703 | 22+<br>21+ | 141,060 | N/A |

| | | | | | | |
|---|---|---|---|---|---|---|
| | | | 7053.8870 | 20+ | | |
| | | | 7425.1089 | 19+ | | |
| | | | 7837.7589 | 18+ | | |
| ACBI3 | CRL2$^{VHL}$ | 141,007 | 6411.4279 | 22+ | 141,064 | N/A |
| | | | 6716.9211 | 21+ | | |
| | | | 7054.3442 | 20+ | | |
| | | | 7425.2074 | 19+ | | |
| | | | 7838.0195 | 18+ | | |
| | CRL2$^{VHL}$-impurity[c] | N/A | 6435.7387 | 22+ | 141,577 | 513[c] |
| | | | 6742.2366 | 21+ | | |
| | | | 7079.7158 | 20+ | | |
| | | | 7452.6289 | 19+ | | |
| | | | 7866.6185 | 18+ | | |
| | CRL2$^{VHL}$-ACBI3 | 142,026.3 | 6458.0264 | 22+ | 142,078 | 1,014 |
| | | | 6766.2734 | 21+ | | |
| | | | 7104.8006 | 20+ | | |
| | | | 7478.9288 | 19+ | | |
| | | | 7894.5842 | 18+ | | |
| *cis*-ACBI3 | CRL2$^{VHL}$ | 141,007 | 6412.8701 | 22+ | 141,080 | N/A |
| | | | 6718.8696 | 21+ | | |
| | | | 7054.8233 | 20+ | | |
| | | | 7426.1097 | 19+ | | |
| | | | 7838.3417 | 18+ | | |

[a]MW deconvoluted in UniDec,[2] to nearest whole Da and with relative intensity >10%. [b]Additional mass corresponds to the binding of either a $Na^+$ (23 Da) or $Mg^{2+}$ (24 Da) adduct. [c]Binding of an ~515 Da impurity in ACBI3 detected. N/A: not applicable as no ligand binding observed.

**Supporting Table S10.** Expected MW of KRAS protein complexes (based on corrected MW).

| Protein | Corrected MW (Da)[a] |
|---|---|
| KRAS | 19,882[b] |
| VCB | 41,373[c] |
| CRL2$^{VHL}$ | 141,007[b] |
| UBE2R1(C93K) | 26,631[c] |
| UBE2R1(C93K)-Ub | 36,247[b] |
| **Compounds/PROTACs** | **MW (Da)** |
| ACBI3 | 1019.3 |
| *cis*-ACBI3 | 1019.3 |
| **Protein Complexes** | **Expected MW (Da)[d]** |
| KRAS-VCB[e] | 61,255 |
| KRAS-ACBI3-VCB[f] | 62,274 |
| KRAS-CRL2$^{VHL}$[e] | 160,889 |
| KRAS-ACBI3-CRL2$^{VHL}$[f] | 161,908 |
| CRL2$^{VHL}$-UBE2R1(C93K)-Ub | 177,254 |
| KRAS-CRL2$^{VHL}$-UBE2R1(C93K)-Ub[e] | 197,136 |
| KRAS-ACBI3-CRL2$^{VHL}$-UBE2R1(C93K)-Ub[f] | 198,155 |

[a]Corrected MW is based on the nMS analysis for protein components (**Supporting Figure S1** and **S7**) and are used for data analysis throughout the rest of the study. [b]MW as determined from nMS visibility studies (**Supporting Figure S1** and **S7**) due to differences with that expected from protein sequence. [c]MW as determined from protein sequence (corresponds with MW as measured by nMS, **Supporting Figure S1**). [d]Expected MWs of protein complexes of interest for this study, calculated using the corrected MWs for individual protein components (rounded to nearest Da). [e]Complexes without PROTAC and not necessarily expected to form, but MWs provided to compare with spectra to confirm presence or absence. [f]Complexes containing *cis*-ACBI3 are not expected to form, however, due to the MW of *cis*-ACBI3, any such complexes would have the same MW as the ACBI3-containing complexes.

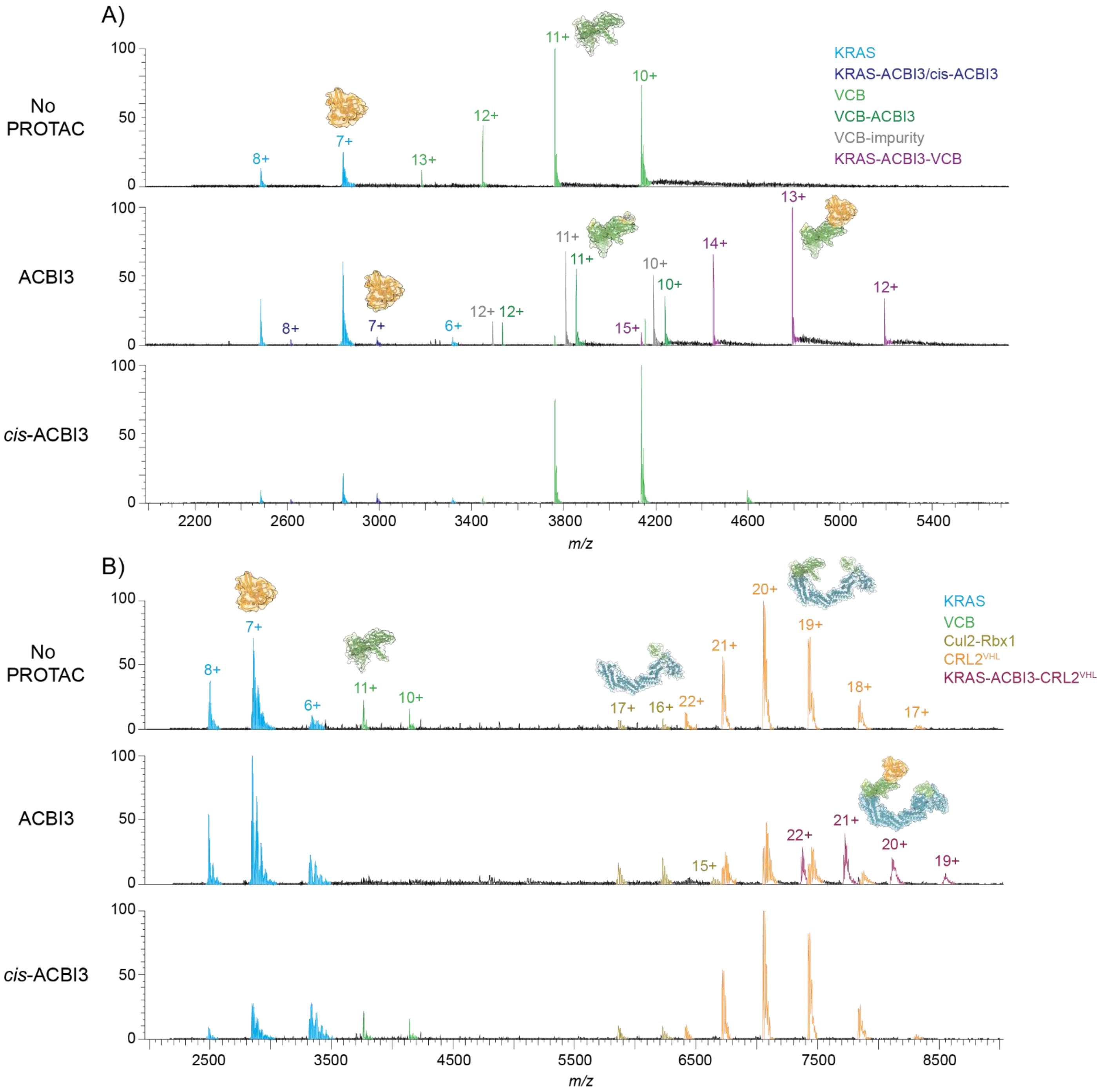

**Supporting Figure S8.** nMS analysis of ternary complex POI-PROTAC-E3 formation between KRAS (1 µM, 1 equiv), ACBI3 (1.9 µM, 1.9 equiv) and A) VCB (1 µM, 1 equiv), or B) CRL2$^{VHL}$ (1 µM, 1 equiv) (150 mM $NH_4OAc$, 1% DMSO). Mass spectra are raw data with charge state and species identity annotated. Top panel is no PROTAC control (1% DMSO), middle panel is with addition of ACBI3, and lower panel is with addition of negative control *cis*-ACBI3 (1.9 equiv). Observed *m/z* values and MWs are in **Supporting Information**, **Table S11**.

**Supporting Table S11.** Observed *m/z* values and measured MWs for samples comprising KRAS and either VCB or CRL2$^{VHL}$ with ACBI3 (MW 1019.3 Da) and *cis*-ACBI3 (MW 1019.3 Da). Raw mass spectra are shown in **Supporting Figure S8**. Binary complexes – blue text, ternary complexes – red text.

| Sample POI | E3 | Compound | Observed Species | Expected MW (Da) | Observed *m/z* | Charge State (*z*) | Deconvoluted MW (Da)[a] |
|---|---|---|---|---|---|---|---|
| KRAS | VCB | - | KRAS | 19,882.0 | 2486.1421 | 8+ | 19,882 |
| | | | | | 2841.2028 | 7+ | |
| | | | VCB | 41,373.4 | 3183.4138 | 13+ | 41,372 |
| | | | | | 3448.6803 | 12+ | |
| | | | | | 3762.1409 | 11+ | |
| | | | | | 4138.2725 | 10+ | |
| | | ACBI3 | KRAS | 19,882.0 | 2486.1345 | 8+ | 19,882 |
| | | | | | 2841.1949 | 7+ | |
| | | | | | 3318.3043 | 6+ | |
| | | | KRAS-ACBI3 | 20,901.3 | 2616.3069 | 8+ | [b] |
| | | | | | 2990.0156 | 7+ | |
| | | | VCB | 41,373.4 | 3761.9889 | 11+ | [b] |
| | | | | | 4138.2183 | 10+ | |
| | | | VCB-impurity[c] | N/A | 3222.9428 | 13+ | 41,887[c] |
| | | | | | 3491.4963 | 12+ | |
| | | | | | 3808.9114 | 11+ | |
| | | | | | 4189.6723 | 10+ | |
| | | | VCB-ACBI3 | 42,392.7 | 3261.9083 | 13+ | 42,391 |
| | | | | | 3533.5278 | 12+ | |
| | | | | | 3854.7394 | 11+ | |
| | | | | | 4240.1162 | 10+ | |
| | | | KRAS-ACBI3-VCB | 62,274.7 | 4154.0189 | 15+ | 62,297[d] |
| | | | | | 4450.77 | 14+ | |
| | | | | | 4793.0842 | 13+ | |
| | | | | | 5192.4958 | 12+ | |
| | | *cis*-ACBI3 | KRAS | 19,882.0 | 2486.1345 | 8+ | 19,904[d] |
| | | | | | 2841.1949 | 7+ | |
| | | | | | 3318.322 | 6+ | |
| | | | KRAS-*cis*-ACBI3 | 20,901.3 | 2616.3069 | 8+ | 20,924[d] |
| | | | | | 2990.0156 | 7+ | |
| | | | VCB | 41,373.4 | 3448.694 | 12+ | 41,372 |
| | | | | | 3762.1354 | 11+ | |
| | | | | | 4138.2702 | 10+ | |
| | | | | | 4597.9777 | 9+ | |
| | CRL2$^{VHL}$ | - | KRAS | 19,882.0 | 2486.4315 | 8+ | 19,939[d] |
| | | | | | 2841.4149 | 7+ | |
| | | | | | 3324.3709 | 6+ | |
| | | | VCB | 41,373.4 | 2762.2050 | 11+ | 41,372 |

| | | | | | |
|---|---|---|---|---|---|
| | | | 4138.3487 | 10+ | |
| | Cul2-Rbx1 | 99,458.2 | 5862.8148 | 17+ | 99,644 |
| | | | 6227.2146 | 16+ | |
| | $CRL2^{VHL}$ | 141,007 | 6412.7951 | 22+ | 141,078 |
| | | | 6718.7686 | 21+ | |
| | | | 7054.806 | 20+ | |
| | | | 7426.3355 | 19+ | |
| | | | 7838.8268 | 18+ | |
| | | | 8300.1577 | 17+ | |
| ACBI3 | KRAS | 19,882.0 | 2486.4035 | 8+ | 19,927[d] |
| | | | 2841.48 | 7+ | |
| | | | 3315.0734 | 6+ | |
| | Cul2-Rbx1 | 99,458.2 | 5860.0158 | 17+ | 99,607 |
| | | | 6226.3478 | 16+ | |
| | | | 6641.8889 | 15+ | |
| | $CRL2^{VHL}$ | 141,007.0 | 6717.7733 | 21+ | 141,048 |
| | | | 7054.015 | 20+ | |
| | | | 7424.5654 | 19+ | |
| | | | 7837.5147 | 18+ | |
| | KRAS-ACBI3-$CRL2^{VHL}$ | 161,908.3 | 7364.7186 | 22+ | 162,251 |
| | | | 7716.3797 | 21+ | |
| | | | 8102.8139 | 20+ | |
| | | | 8543.4401 | 19+ | |
| *cis*-ACBI3 | KRAS | 19,882.0 | 2486.3181 | 8+ | 20,002[d] |
| | | | 2841.5444 | 7+ | |
| | | | 3318.497 | 6+ | |
| | VCB | 41,373.4 | 3762.205 | 11+ | 41,373 |
| | | | 4138.3487 | 10+ | |
| | Cul2-Rbx1 | 99,458.2 | 5859.4455 | 17+ | 99,595 |
| | | | 6226.1817 | 16+ | |
| | $CRL2^{VHL}$ | 141,007 | 6411.4829 | 22+ | 141,059 |
| | | | 6717.0906 | 21+ | |
| | | | 7053.6539 | 20+ | |
| | | | 7425.2275 | 19+ | |
| | | | 7837.7281 | 18+ | |
| | | | 8299.9313 | 17+ | |

[a]MW deconvoluted in UniDec,[2] to nearest whole Da and with relative intensity >5%. Increased mass compared to expected M corresponds to adducts such as $Na^+$ or $Mg^{2+}$. [b]Species not detected during deconvolution but evident in raw spectrum. [c]Binding of an ~515 Da impurity present in the ACBI3 sample observed. N/A: not applicable as exact impurity MW unknown.

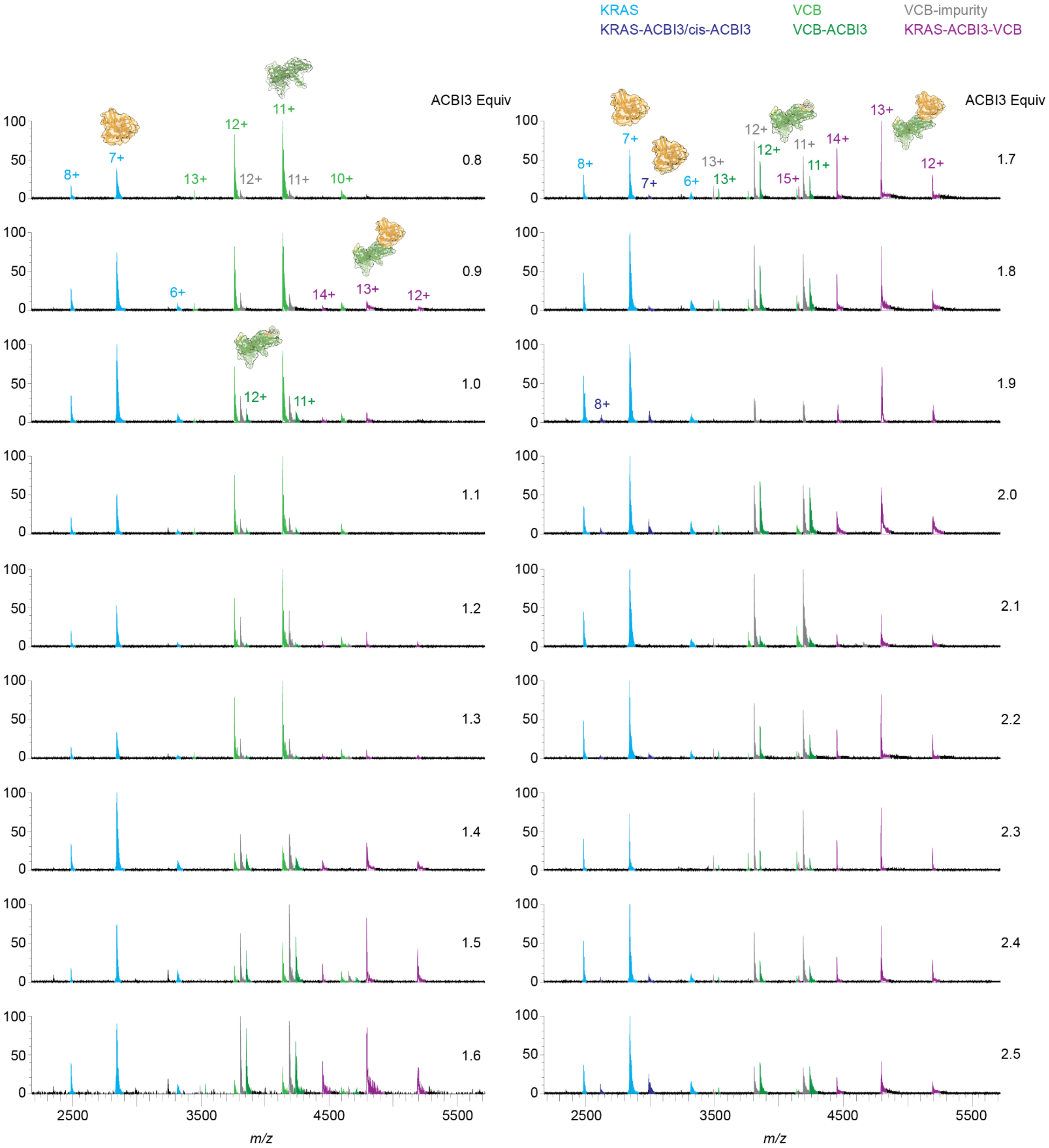

**Supporting Figure S9.** nMS titration of PROTAC ACBI3 (0.8-2.5 equiv, 0.1 increments) against POI KRAS and E3 VCB (present at equimolar concentrations, 1 µM each) to determine the optimal ACBI3 equiv for maximal ternary complex formation.

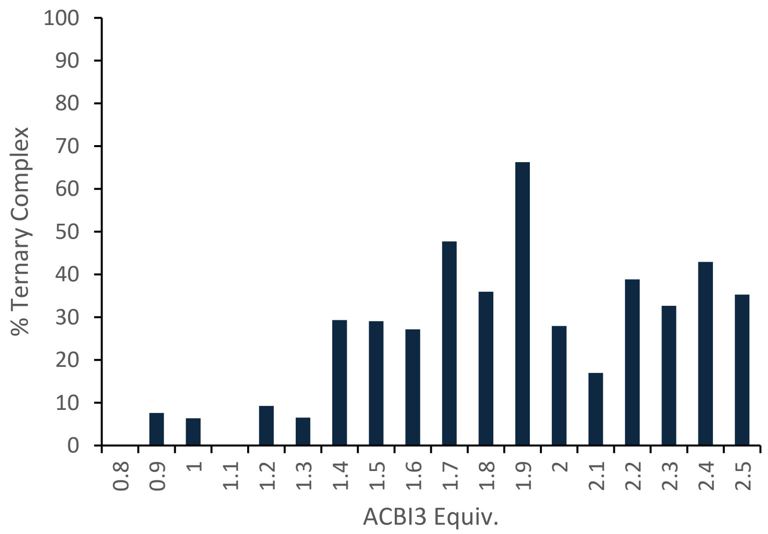

**Supporting Figure S10.** Quantification of ternary complex formation between KRAS and VCB (present at equimolar concentrations, 1 µM each) with ACBI3 titrated (0.8-2.5 equiv, 0.1 increments) as measured by nMS. The % ternary complex formed was calculated from the deconvoluted data of mass spectra in **Supporting Figure S9** as the ratio of VCB recruited into the ternary complex (i.e. sum of the intensity of the deconvoluted ternary complex peak divided by the sum of the intensities of the deconvoluted peaks for unbound VCB, impurity-bound VCB, ACBI3-bound VCB and ternary complex).

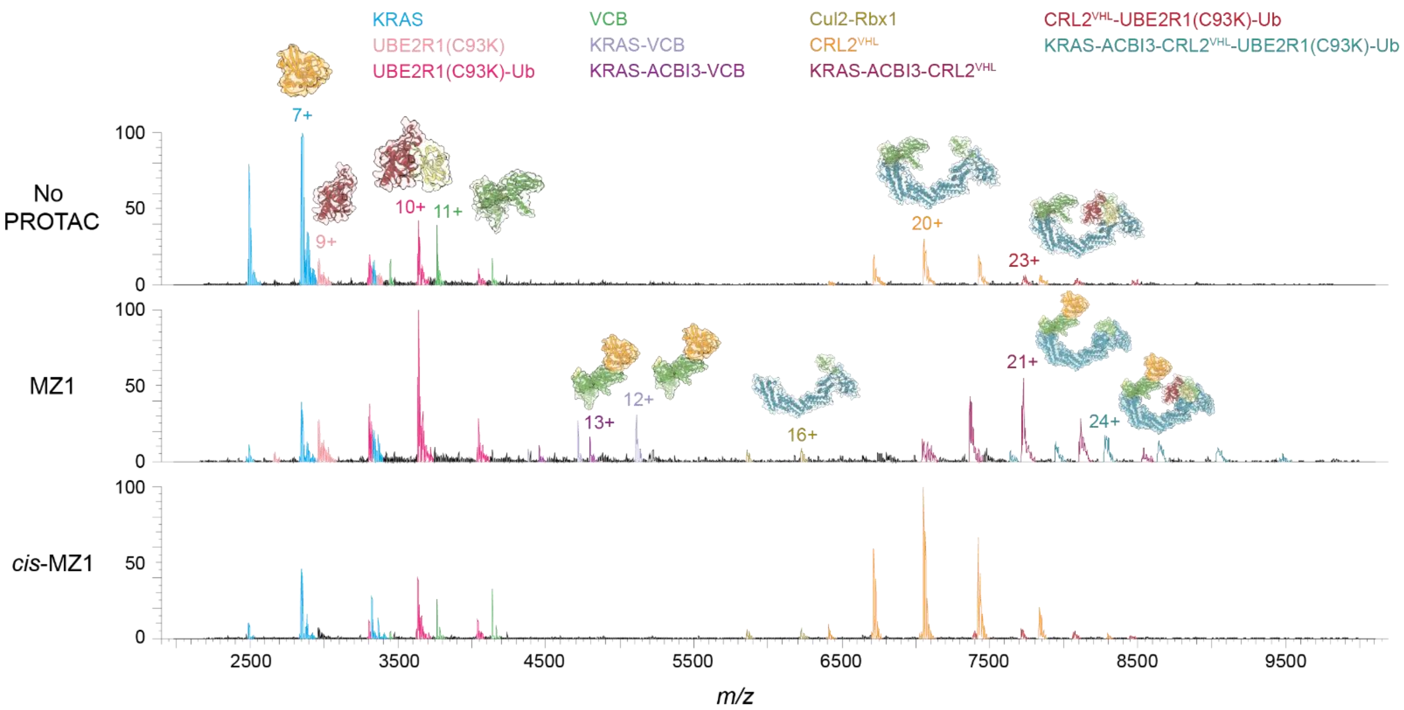

**Supporting Figure S11.** Raw data nMS spectra showing PROTAC mediated formation of the full KRAS-ACBI3-CRL2$^{VHL}$-UBE2R1(C93K)-Ub complex. The peaks for each observed species that were present at >3% relative intensity following deconvolution are labelled via the indicated colour coding, and the charge state for the most intense peak for each observed species is also labelled. Corresponding deconvoluted nMS are in manuscript **Figure 6D-F** and observed *m/z* values and MWs are provided in **Supporting Table S12**.

**Supporting Table S12.** Observed *m/z* values and measured MWs for the full POI-PROTAC-E3-E2-Ub degradation complex with KRAS as the POI. Raw and deconvoluted spectra are shown in **Supporting Figure S10** and **Figure 6D-F**, respectively). Binary complexes – blue text, ternary complexes – red text, full complex in green text.

| Sample Proteins | Compound | Protein Species | Expected MW (Da) | Observed *m/z* | Charge State (*z*) | Deconvoluted MW (Da)[a] |
|---|---|---|---|---|---|---|
| KRAS, CRL2$^{VHL}$, UBE2R1(C93K)-Ub | - | KRAS | 19,882.0 | 2486.2542<br>2841.3606 | 8+<br>7+ | 19,904 |
| | | UBE2R1(C93K) | 26,630.6 | 2959.9899<br>3330.6674 | 9+<br>8+ | 26,632 |
| | | UBE2R1(C93K)-Ub | 36,247.0 | 3301.6323<br>3631.6572<br>4035.0980 | 11+<br>10+<br>9+ | 36,307 |
| | | VHL-EloB-EloC | 41,373.4 | 3448.6136<br>3762.2210<br>4138.3816 | 12+<br>11+<br>10+ | 41,373 |
| | | CRL2$^{VHL}$ | 141,007.0 | 6413.8685<br>6717.8919<br>7053.7534<br>7425.6475<br>7838.8709 | 22+<br>21+<br>20+<br>19+<br>18+ | 141,060 |
| | | CRL2$^{VHL}$-UBE2R1(C93K)-Ub | 177,254.0 | 7728.0971<br>8080.9648<br>8465.0562 | 23+<br>22+<br>21+ | 177,734 |
| | ACBI3 | KRAS | 19,882.0 | 2493.0915<br>2849.0872 | 8+<br>7+ | 19,936 |
| | | UBE2R1(C93K) | 26,630.6 | 2663.9497<br>2959.9316 | 10+<br>9+ | 26,630 |
| | | UBE2R1(C93K)-Ub | 36,247.0 | 3301.6092<br>3631.6854<br>4035.2016 | 11+<br>10+<br>9+ | 36,307 |
| | | KRAS-VCB | 61,255.4 | 4380.4830<br>4717.5760<br>5110.8157 | 14+<br>13+<br>12+ | 61,317 |
| | | KRAS-ACBI3-VCB | 62,247.7 | 4453.4398<br>4796.2024 | 14+<br>13+ | 62,337 |
| | | Cul2-Rbx1 | 99,458.2 | 5859.5129<br>6225.5701 | 17+<br>16+ | 99,593 |
| | | KRAS-ACBI3-CRL2$^{VHL}$ | 161,908.3 | 7044.6393<br>7365.2088<br>7716.7247 | 23+<br>22+<br>21+ | 162,023 |

| | | | | | |
|---|---|---|---|---|---|
| | | | 8103.2198 | 20+ | |
| | | | 8530.3381 | 19+ | |
| | KRAS-ACBI3-CRL2$^{VHL}$-UBE2R1(C93K)-Ub | 198,155.3 | 7638.5919 | 26+ | 198,644 |
| | | | 7944.2201 | 25+ | |
| | | | 8276.8724 | 24+ | |
| | | | 8642.0228 | 23+ | |
| | | | 9034.5203 | 22+ | |
| | | | 9459.2066 | 21+ | |
| *cis*-ACBI3 | KRAS | 19,882.0 | 2486.3380 | 8+ | 19,882 |
| | | | 2841.3167 | 7+ | |
| | | | 3323.8015 | 6+ | |
| | UBE2R1(C93K)-Ub | 36,247.0 | 3301.6776 | 11+ | 36,307 |
| | | | 3631.7890 | 10+ | |
| | | | 4035.2441 | 9+ | |
| | VHL-EloB-EloC | 41,373.4 | 3448.5657 | 12+ | 41,374 |
| | | | 3762.2138 | 11+ | |
| | | | 4138.3817 | 10+ | |
| | Cul2-Rbx1 | 99,458.2 | 5859.2432 | 17+ | 99,591 |
| | | | 6225.5027 | 16+ | |
| | CRL2$^{VHL}$ | 141,007.0 | 6409.1202 | 22+ | 140,991 |
| | | | 6714.6777 | 21+ | |
| | | | 7050.4881 | 20+ | |
| | | | 7421.7902 | 19+ | |
| | | | 7834.6201 | 18+ | |
| | | | 8296.4481 | 17+ | |
| | CRL2$^{VHL}$-UBE2R1(C93K)-Ub | 177,254.0 | 7391.1594 | 24+ | 177,373 |
| | | | 7712.3778 | 23+ | |
| | | | 8063.4043 | 22+ | |
| | | | 8450.1208 | 21+ | |

[a]MW deconvoluted in UniDec,[2] to nearest whole Da and with relative intensity >3%. Increased mass compared to expected MW corresponds to adducts such as $Na^+$ or $Mg^{2+}$.

**Recombinant production of CRL2$^{VHL}$ protein.**

50 µL of DH10Bac competent *E. coli* cells were thawed on ice. 1 µL (600 ng) plasmid DNA[1] was added and incubated on ice for 30 min. The cells were heat shocked at 42 °C for 90 seconds before incubating on ice for 5 min. 500 µL Luria Broth (LB) media was added under laminar flow and the cells were incubated at 37 °C for 5 h. 90 uL of the cell suspension was plated onto insect cell medium agar plates (containing 50 µg/mL kanamycin, 7 µg/mL gentamicin, 10 µg/mL tetracycline, 40 µg/mL IPTG and 100 µg/mL X-Gal) under laminar flow and incubated at 37 °C for 48 h.

5 mL of insect cell media (containing 50 µg/mL kanamycin, 7 µg/mL gentamicin, 10 µg/mL tetracycline) was inoculated with single white colonies. The freshly inoculated media was streaked onto new insect cell medium agar plates (containing 50 µg/mL kanamycin, 7 µg/mL gentamicin, 10 µg/mL tetracycline, 40 µg/mL IPTG and 100 µg/mL X-Gal) to discard false positives. The cultures were incubated at 37 °C with shaking at 180 rpm overnight. The plates were incubated at 37 °C for 48 h. The cells were pelleted and the bacmid was isolated through mini-prep. The bacmids were stored at 4 °C until transfection, before longer term storage at -20 °C. Successful recombination was verified by PCR.

*Spodoptera frugiperda 9 (Sf9)* insect cells were grown in suspension culture, using Gibco Sf-900™ II SFM media (ThermoFisher) supplemented with Gibco Antibiotic-Antimycotic (ThermoFisher). The cells were incubated at 27 °C with 130 rpm shaking, protected from light. The cells were split every 72 to 96 h to a density of $1.0 \times 10^6$ cells/mL. Cell count, viability and cell diameter were monitored using a Luna fl Cell Counter. The cell viability was maintained above 90% and the cell diameter of uninfected cells was usually 13-17 µm.

Sf9 insect cells were seeded at $1.0 \times 10^6$ cells/mL, one day prior to transfection. Once cells had reached $1.5 \times 10^6$ cells/mL, 5 mL of cells were seeded at $0.8 \times 10^6$ cells/mL into T25 flasks. The cells were allowed to attach for 15 min at room temperature. The media was aspirated and replaced with 3 mL of fresh Gibco Sf-900™ II SFM media. To a solution of 200 µL of Gibco Sf-900™ II SFM media and 8 µL of Cellfectin II (ThermoFisher), was added 3 µg of the bacmid DNA. The resulting mixture was mixed and incubated at room temperature for 30 min. 300 µL Gibco Sf-900™ II SFM media was added, and the transfection mixture was added to the cells dropwise. The culture was gently mixed, then incubated at 27 °C for 6 h. The media was aspirated and replaced with 5.5 mL of fresh Gibco Sf-900™ II SFM media supplemented with Gibco Antibiotic-Antimycotic. The culture was incubated for 6 days at 27 °C. The supernatant was collected and centrifuged at 1500 × g at room temperature for 5 min. The supernatant (P0 baculovirus stock) was collected, transferred to fresh flasks, and stored protected from light at 4 °C. For virus amplification, to a freshly seeded 300 mL culture of Sf9 cells at $1.0 \times 10^6$ cells/mL was added 1 mL of P0 baculovirus stock. The culture was incubated for 4 days at 27 °C with shaking at 130 rpm, protected from light. The culture was centrifuged at

800 × g at room temperature for 5 min. The supernatant (P1 baculovirus stock) was collected, transferred to fresh flasks, and stored protected from light at 4 °C.

Sf9 cells were seeded at a density of 1.0 × $10^6$ cells/mL. After 24 h of incubation, the cells were infected with P1 baculovirus stock in a ratio of 1:50, 1:75 or 1:100. The culture was incubated for 72 hat 27 °C with shaking at 130 rpm, protected from light. The cells were harvested by centrifugation at 4000 rpm, 4 °C for 30 min. The cell pellets were resuspended in 50 mM HEPES, 250 mM NaCl, 2 mM TCEP and 0.2% (v/v) Triton-X, pH 8.0, flash frozen at stored at -80 °C until purification.

To the thawed and resuspended cell pellets was added cOmpleteTM EDTA-free protease inhibitor cocktail (Roche), 5 mM $MgCl_2$ and 10 µg/mL DNAse I. The mixture was incubated for 15 min at room temperature, and the cells were lysed at 15 kpsi on a Constant Cells cell disruptor. The protein-containing supernatant was clarified by centrifugation at 22,000 rpm for 20 min at 4 °C. Ampicillin-Sepharose resin was equilibrated in water, and the lysate was incubated with the resin for 50 min at room temperature. The resin was spun at 7000 rpm for 2 min, washed with 50 mM HEPES, 250 mM NaCl, 2 mM TCEP and 0.2% (v/v) Triton-X, pH 8.0, and was left in the same buffer. TEV protease was added along with an excess of purified recombinant VCB protein, and the protein was cleaved at room temperature for 2.5 h. The mixture was filtered, and the filtrate was purified on a 10/300 GL Superdex 200 Increase prepacked column (Cytiva) eluting in 20 mM HEPES pH 8.0, 100 mM NaCl, 5% (v/v) glycerol, 2 mM TCEP.

**Recombinant production of KRAS protein.**

KRAS (residues 1 to 169) wild type was expressed from a pET-s8a(+) expression vector and purified in *E. coli* BL21(DE3). An overnight culture was grown in LB supplemented with 50 µg/mL kanamycin at 37 °C, 180 rpm. To each 1L of fresh LB media (containing 50 µg/mL kanamycin), 10 mL overnight culture was added and further grown at 37 °C, 180 rpm. Cultures were grown until an OD600 of 0.8 was reached, after which cells were induced with 250 µM IPTG and the temperature was reduced to 18 °C. KRAS was expressed overnight and harvested the next morning by centrifugation at 4000 rpm, for 40 min, 4 °C using a Beckman Coulter Avanti J-HC equipped with a JS-4.2 rotor. The pellets were stored at -20 °C in approximately 200 mL of Buffer A (20 mM Tris, pH 8.0, 500 mM NaCl, 10 mM $MgCl_2$, 1 mM TCEP), supplemented with 2 tablets of cOmplete Protease Inhibitor Cocktail tablets (Roche) and 50 µL 140 µM benzonase. The pellets were thawed to room temperature prior to cell-disruption using a One Shot Cell Disrupter (Constant Systems) operating at 30 kpsi, 4 °C. The cell-debris was separated from the lysate by centrifugation at 25 000 rpm for 40 min at 4 °C, using a Beckman Coulter Avanti J-26S XP with a JA-25.50 rotor. The cleared lysate was syringe filtered through a 0.45 µm filter before running it over a Ni-affinity HisTrap HP column (Cytiva). The protein was eluted using a 10 to 300 mM imidazole gradient over 10 column volumes (CV) run at 5 mL/min in Buffer A. Fractions containing KRAS were combined and subjected to TEV protease cleavage (300 µL of

10 mg/mL TEV protease) while dialysing to Buffer A at 4 °C overnight. KRAS was further purified by running a reverse HisTrap using a 10 to 300 mM imidazole gradient over 10 CV run at 5 mL/min in Buffer A. Fractions containing KRAS were pooled and concentrated to a volume of 1.8 mL, after which it was further purified using gel filtration. A Cytiva HiLoad Superdex 75 pg 16/600 column was used at 0.35 mL/min, 4 °C using Buffer B (20 mM Tris, pH 7.5, 10 mM NaCl, 2 mM $MgCl_2$, 1 mM TCEP). Fractions containing pure KRAS were combined and concentrated to about 3 mg/mL. To the pure KRAS solution 0.5 mg of guanosine 5′-diphosphate (GDP) per mg of protein was added and incubated at 4 °C for 5.5 h. After incubation with GDP the buffer was exchanged to Buffer B without GDP and concentrated to 2 mM. 31 aliquots of 20 µL were flash frozen in liquid nitrogen and stored at -80 °C.

**Supporting Table S13.** UHMR MS parameters employed for the methods used to characterise proteins and/or complexes. Typically, 1-2 mins of acquired data were averaged per spectrum. nMS data were analysed directly using Themo Fisher Scientific FreeStyle 1.8 SP2 QF1 and deconvoluted in UniDec.[2]

| **Parameter** | **BRD4$^{BD2}$ Binary Binding** | **KRAS Binary Binding** | **VCB Binary Binding** | **VCB Ternary Complexes** | **CRL2$^{VHL}$ Binary Binding and CRL2$^{VHL}$ Ternary Complexes** | **E3-E2 Interactions and BRD4$^{BD2}$-PROTAC-E3-E2-Ub Complexes** | **KRAS-PROTAC-E3-E2-Ub Complexes** |
|---|---|---|---|---|---|---|---|
| Polarity | Positive | Positive | Positive | Positive | Positive | Positive | Postitive |
| nESI Source | NanoMate | NanoMate | NanoMate | NanoMate | Static | Static | Static |
| Spray voltage (V) | 1.7 | 1.7 | 1.7 | 1.7 | 0.8-1.1 | 0.8-1.4 | 0.8-1.2 |
| NM pressure (psi) | 1.0 | 1.0 | 1.0 | 1.0 | N/A | N/A | N/A |
| Capillary temperature (°C) | 275 | 275 | 275 | 230 | 240 | 275 | 240 |
| S-lens RF level | 200 | 200 | 200 | 200 | 200 | 200 | 200 |
| In-source trapping desolvation voltage (V) | -5 | 0 | -5 | -10 | -60 | -40 | -60 |
| Source DC offset (V) | 5 | 21 | 5 | 21 | 21 | 21 | 21 |
| Injection flatapole DC (V) | 5 | 5 | 5 | 5 | 5 | 5 | 5 |
| Inter flatapole lens (V) | 4 | 4 | 4 | 4 | 4 | 4 | 4 |
| Bent flatapole DC (V) | 2 | 2 | 2 | 2 | 2 | 2 | 2 |
| Transfer multipole DC (V) | 0 | 0 | 0 | 0 | 0 | 0 | 0 |
| C-trap entrance lens (V) | 1.8 | 1.8 | 1.8 | 1.8 | 1.8 | 1.8 | 1.8 |

| | | | | | | | |
|---|---|---|---|---|---|---|---|
| Ion transfer target *m/z* | High *m/z* | High *m/z* | High *m/z* | High *m/z* | High *m/z* | High *m/z* | High *m/z* |
| Trapping gas pressure | 1.0 | 3.0 | 1.0 | 1.0 | 4.0 | 4.0 | 4.0 |
| Extended trapping | 1.0 | 1.0 | 1.0 | 1.0 | 10.0 | 10.0 | 10.0 |
| Detector *m/z* optimisation | Low *m/z* | Low *m/z* | Low *m/z* | Low *m/z* | Low *m/z* | Low *m/z* | Low *m/z* |
| Scan range *m/z* | 1,500-6,000 | 1,500-6,000 | 1,500-6,000 | 1,500-8,000 | 2,000-10,000 | 2,000-10,000 | 2,000-10,000 |
| Resolution | 200,000 | 100,000 | 25,000 | 12,500 | 6,250 | 6,250 | 6,250 |
| Microscans | 5 | 5 | 5 | 5 | 10 | 10 | 10 |
| Manual inject time | 5 | 100 | 100 | 100 | 200 | 200 | 200 |
| Averaging | 5 | 5 | 5 | 5 | 5 | 10 | 5 |
| **UniDec Processing** | | | | | | | |
| *m/z* range | 1,500-6,000 | 1,500-6,000 | 1,500-8,000 | 1,500-8,000 | 2,000-10,000 | 2,000-10,000 | 2,000-10,000 |
| Charge state range | 4-7 | 3-12 | 4-15 | 3-40 | 3-30 or 40 | 3-40 | 3-50 |
| MW range (Da) | 10,000-100,000 | 5,000-30,000 | 10,000-100,000 | 5,000-200,000 | 5,000-200,000 | 10,000-200,000 or 250,000 | 5,000-250,000 |
| Sample mass every x Da | 1 | 1 | 1 | 1 | 1 | 1 | 1 |
| Peak detection range (Da) | 500 | 500 | 500 | 500 | 500 | 500 | 500 |
| Peak detection threshold (Da) | 0.1 | 0.05 | 0.1 | 0.05 | 0.05 or 0.1 | 0.05 (lowered to 0.03 where required) | 0.05 (lowered to 0.03 where required) |
